# Evaluation of established methods for DNA extraction and primer pairs targeting 16S rRNA gene for bacterial microbiome profiling of olive xylem sap

**DOI:** 10.1101/2020.12.11.420356

**Authors:** Carmen Haro, Manuel Anguita-Maeso, Madis Metsis, Juan A. Navas-Cortés, Blanca B. Landa

## Abstract

Next Generation Sequencing has revolutionized our ability to investigate the microbiota composition of diverse and complex environments. However, a number of factors can affect the accuracy of microbial community assessment, such as the DNA extraction method, the hypervariable region of 16S rRNA gene targeted or the PCR primers used for amplification. The aim of this study was to assess the influence of commercially available DNA extraction kits and different primer pairs to provide a nonbiased vision of the composition of bacterial communities present in olive xylem sap. For that purpose, branches from ‘Picual’ and ‘Arbequina’ olive cultivars were used for xylem sap extraction using a Scholander chamber device. The DNA extraction protocol significantly affected xylem sap bacterial community assessment. That resulted in significant differences in alpha (Richness) and beta diversity (UNIFRAC distances) metrics among DNA extraction protocols, with the 12 DNA extraction kits evaluated being clustered in four groups behaving differently. Although the core number of taxa detected by all DNA extraction kits included four phyla, seven classes, 12 orders, and 16 or 21 families, and 12 or 14 genera when using the Greengenes or Silva database for taxonomic assignation, respectively, some taxa, particularly those identified at low frequency, were detected by some DNA extraction kits only. The most accurate depiction of a bacterial mock community artificially inoculated on sap samples was generated when using the PowerPlant DNA extraction Kit, the combination of 799F/1193R primers amplifying the hypervariable V5-V7 region and the Silva 132 database for taxonomic assignation. The DESeq2 analysis displayed significant differences among genera abundance between the different PCR primer pairs tested. Thus, *Enterobacter, Granulicatella, Prevotella* and *Brevibacterium* presented a significant higher abundance in all PCR protocols when compared with primer pair 799F/1193R, while the opposite was true for *Pseudomonas* and *Pectobacterium*. The methodological approach followed in this study can be useful to optimize plant-associated microbiome analysis, especially when exploring new plant niches. Some of the DNA extraction kits and PCR primers selected in this study will contribute to better characterize bacterial communities inhabiting within the xylem sap of olives or other woody crop species.

## Introduction

Olive (*Olea europaea* subsp. *europaea*) is a primary element in the agricultural economy of most countries in the Mediterranean Basin, where about 5 million hectares of olive orchards are grown only in European countries. More than half of the cultivated olives worldwide are in Spain, accounting for 70 to 75 % of world production of olive oil and more than one third for table olives (EUROSTAT, https://ec.europa.eu/eurostat). During the last years, the health of the olive groves is being seriously threatened, as consequence of a notable increase, both in extent and in severity, of diseases caused by diverse pathogens, which are capable of adversely affect its growth and production. Among olive diseases, those caused by the vascular plant pathogenic bacterium *Xylella fastidiosa* and the soilborne fungus *Verticillium dahliae* are, without a doubt, the two major global threats for olive production worldwide (Jiménez-Díaz et al., 2011; Saponari et al., 2018; Landa et al., 2019; Anguita-Maeso et al., 2020).

Research on plant-associated microorganisms or plant microbiome has gained importance in the last decade as a key component in the health and productivity of the plant (Berg et al., 2014). Thus, recent studies have shown that the presence of certain endophytic bacteria are capable of modifying the development of diseases in the plants, promoting their growth and protecting them against insects and pathogens (Müller et al., 2015). In addition, they could confer other important benefits for plants, such as greater resistance to stress conditions, alteration in physiological properties, and production of phytohormones and other compounds of biotechnological interest (Porras-Alfaro and Bayman, 2011; Hacquard and Schadt, 2015; Santoyo et al., 2016). Within the plant tissues, xylem vessels are considered ideal niches for microorganism by providing an effective internal pathway for distribution throughout the plant and a continuous source of nutrients (McCully, 2001). However, the nature and role of the xylem microbiome, and its contribution to plant health and crop productivity is still scarce (Anguita-Maeso et al., 2020).

For olive trees, most microbiome studies have focused on determining the microbial composition of its rhizosphere (Mercado-Blanco et al., 2004; Aranda et al., 2011; Berg et al., 2016; Gómez-Lama Cabanás et al., 2018; Lei et al., 2019; Fernández-González et al., 2020) and to which extent these microbial communities can act as potential antagonists of olive pathogens such as *V. dahliae*. Other studies have focused on the effect of abiotic factors (edaphic, climatic and agronomic) on the olive soil- and rhizosphere-associated microbiome (Montes-Borrego et al., 2013; Landa et al., 2014; Caliz et al., 2015). More scarce are the works in which the endosphere olive microbiome has been investigated, and when done a variety of methodological approaches based on next generation sequencing (NGS) technologies were used to analyze the microbiome composition have been diverse (Müller et al., 2015; Fausto et al., 2018; Sofo et al., 2019; Anguita-Maeso et al., 2020). Moreover, it has been demonstrated that the method used for DNA extraction can lead to dramatic di erences in microbial output composition (Henderson et al., 2013; Brooks et al., 2015), which makes validation of DNA extraction methods with a mock microbial community essential to ensure an accurate representation of the microbial communities in the samples under study.

Most of the referred studies analyzing xylem microbiome have been based on amplicon sequencing using ‘universal’ primers targeting the 16S rRNA gene in bacteria as is the most cost-e□ective and facile tool to provide valuable phylogenetic information for the comparison of bacterial diversity in large numbers of samples. However, the lack of standardization procedures among plant microbiome studies can make difficult the comparison of results among them (Stulberg et al., 2016). Both, choice of the hypervariable region of 16S and the primer pair, have been shown to influence the description of microbial diversity (Claesson et al., 2010). Thus, care should be taken in choosing appropriate primer pairs, as limited taxa coverage, over- or underrepresentation of taxa in a specific environment due to biases in primer amplification could occur, that could lead to unreliable results (Claesson et al., 2010; Wasimuddin et al., 2020).

An additional problem when working with plant tissues is the co-amplification of undesirable or non-target sequences from organellar origin (e.g., mitochondria and/or chloroplast DNA) that may represent a major source of ‘contamination’ due to the homology between bacterial 16S rDNA, chloroplast DNA, and mitochondrial DNA. This leads to significant challenges in the selection of appropriate primer pairs to address the study of plant-microbe interactions (Ghyselinck et al., 2013). Several methodologies have been proposed to reduce co-amplification of plant organellar sequences such as reduction of co-extraction of organellar DNA based on differences in methylation density (Feehery et al., 2013), blocking primers and suicide polymerase endonuclease restriction (SuPER) (Green and Minz, 2005), and the use of specific mismatch primers during PCR amplification (Beckers et al., 2016). Among them, the preferred or most used approach is the use of specific mismatch primers, which amplify bacterial 16S rDNA sequences while simultaneously avoiding the amplification of organellar DNA sequences. Thus, several primer pairs have been developed with that purpose revealing different performance depending of the study (e.g., Chelius and Triplett, 2001; Sogin et al., 2006; Walker and Pace, 2007; Beckers et al., 2016; Dos Santos et al., 2017). However, the experimental performance of these mismatch primers, and their efficacy in reducing co-amplification of non-target DNA in different plant species or plant compartments that may differ on organellar input has not been evaluated enough. Consequently, it is essential to evaluate the amplification efficiency and robustness of selected primer pairs in plant-bacteria interaction studies to assess their behavior in different host plants and specific plant compartments, especially on those rarely addressed such as the xylem tissue.

To our knowledge, no study has systematically evaluated the effects of different DNA extraction methods and the choice of primer pairs to conduct studies on xylem sap bacterial microbiome, which is a key information that is still missing. Consequently, the objectives of this work have been: 1) to compare several standard DNA extraction kits and primer pairs targeting different hypervariable regions of the 16S rRNA gene and described as suitable to avoid co-amplification of plant organellar rRNA gene sequences for their efficacy in the description of the structure and diversity of xylem sap bacterial microbiome when coupled with next generation sequencing and bioinformatics tools; 2) to assess whether one of the selected DNA extraction methods combined with different primer pairs could provide an accurate representation of bacterial communities using a mock microbial community standard; and 3) to demonstrate the utility of the selected protocol for assessing differences in xylem bacterial composition of two of the most widely grown olive cultivars in Spain.

## Materials and Methods

### Xylem Sap Collection

Xylem sap extraction from olive branches was performed using a Scholander pressure chamber pressurized with compressed nitrogen and coupled with an external 60-cm long super chamber following the Bollard process and as described by Anguita-Maeso et al. (2020). Shortly, xylem sap was extracted from 30 cm-long, 2-year old branches of adult olive trees, which were approximately 1 cm in diameter in their thicker part, and which were debarked at the external part to avoid contamination by phloem fluids. Pressure was increased progressively up to a maximum of 35 bar (3.5 MPa), with the first few drops of sap being discarded; then the xylem sap was collected into 5-ml Eppendorf tubes kept on ice for 20-30 minutes until a minimum volume of 4.0 ml per branch was obtained. Xylem sap was kept at −80°C prior to DNA extraction. All the processes described above took place under sterile conditions into a flow hood chamber.

### Extraction of microbial DNA from xylem sap

For the initial experiment to evaluate different microbial DNA extraction protocols, a total of 15 olive branches were used from four 8-year old olive trees of cv. ‘Picual’. The xylem sap extracted from all branches was combined into a composite sample, vortexed to homogenize it and then split into 3 ml aliquots. Samples were then centrifuged at 12,000 x g for 20 minutes at 4°C. The supernatant was removed into a new tube and the pellet and the supernatant were stored at −20°C separately.

Xylem sap pellets were used for total DNA extraction using 11 commercial microbial DNA extraction kits following the protocol as indicated by the manufacturer or with slight modifications (**Table 1**). Additionally, the CTAB protocol (2% Hexadecyl trimethyl-ammonium bromide, 0.1 M Tris-HCl pH 8, 20 mM EDTA and 1.4 M NaCl) was included. Two replicates of xylem sap samples were used for each protocol. Before starting each protocol, the pellet was resuspended in the corresponding extraction buffer of each protocol by vortexing briefly. DNA was eluted in a final volume of 50 μL of ultrapure, filtered-sterilized distilled water. For each protocol and replicated sample, the following parameters were determined: 1) DNA yield by using a NanoDrop^®^156 ND-1000 UV-Vis spectrophotometer (Thermo Fisher Scientific, Inc., Waltham, MA, USA), 2) the PCR yield using PCR primers 967F / 1193R and agarose gel electrophoresis (see below), 3) time of processing and 4) cost (**Table 1**).

**Table 1.**
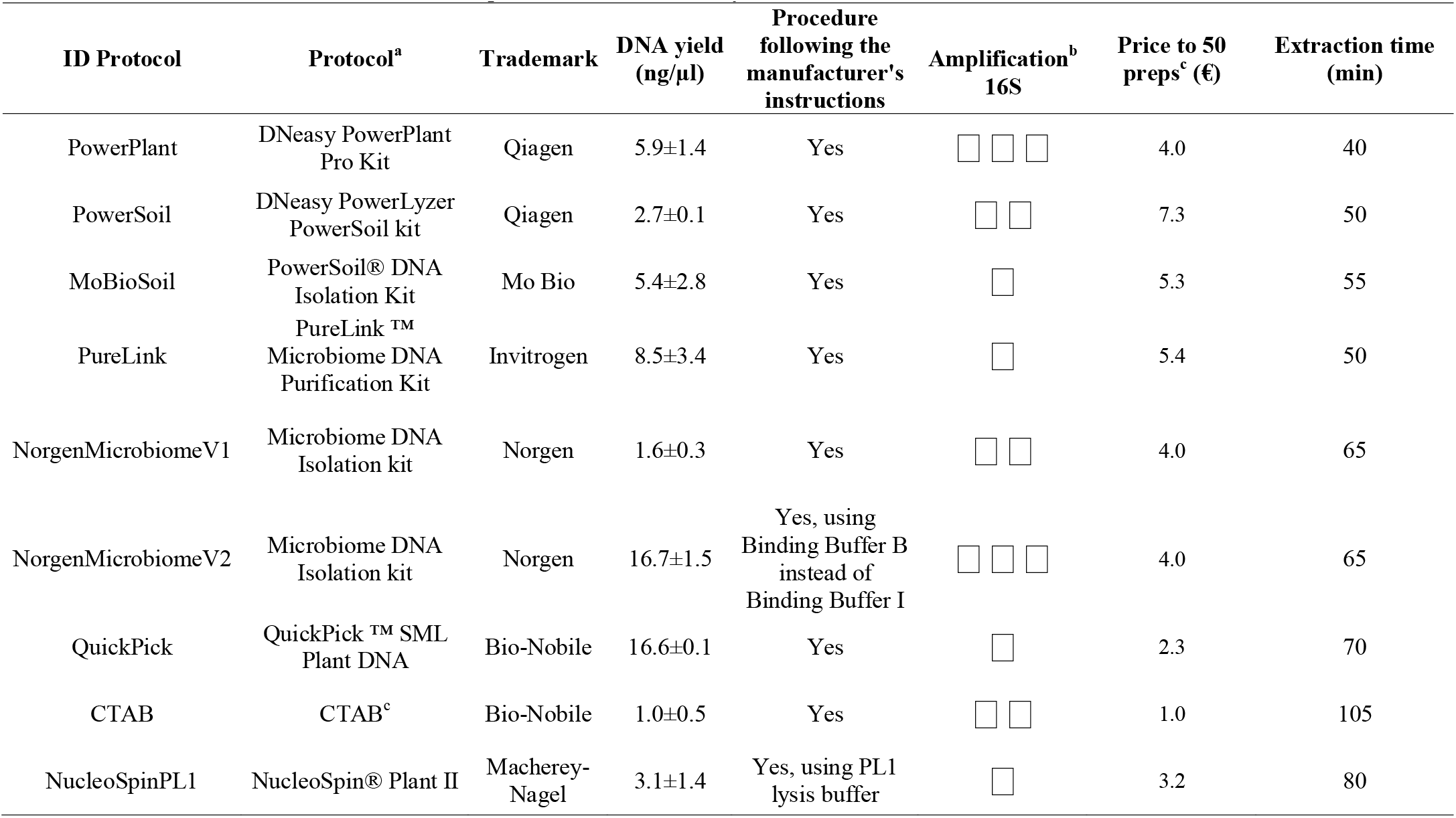
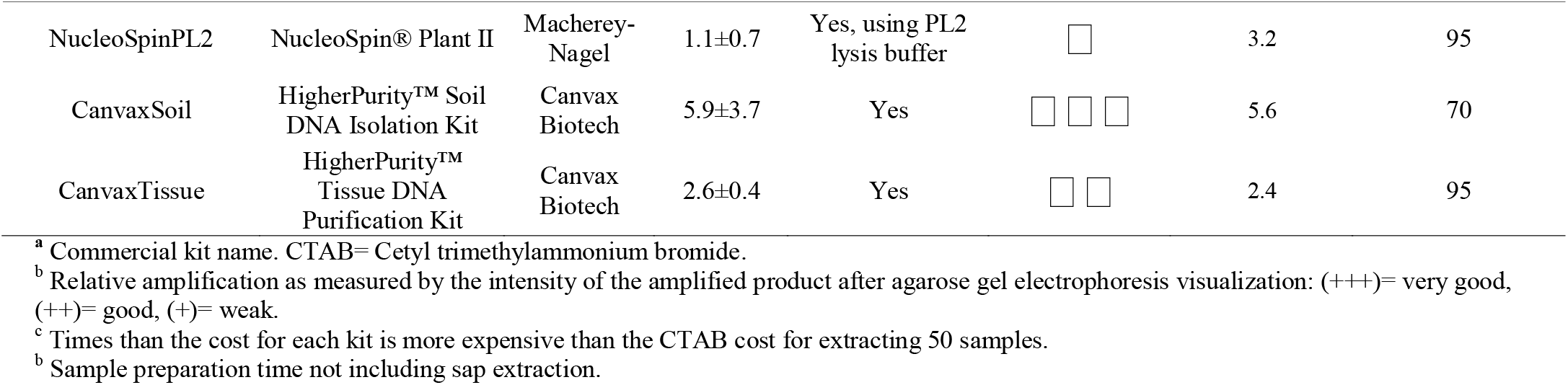
Characteristics of the DNA extraction protocols used in the study.

### Validation of a DNA extraction protocol and PCR primers

Microbial composition profiling techniques powered by Next-Generation Sequencing can suffer from significant bias at different steps from bacterial sampling to bioinformatics analysis workflows. A mock standard microbial community was used to validate which of the four primer pairs combinations could provide the most accurate representation of the microbial communities present in the xylem sap and to which extent the selection of the reference database for taxonomic assignation could also influence results. The ZymoBIOMICS microbial standard (Zymo Research) was used as a mock standard microbial community. This is a well-defined, accurately characterized mock community consisting of three easy-to-lyse bacteria, five tough to-lyse bacteria, and two tough-to-lyse yeasts. Its theoretical bacterial composition according to the manufacturer based on 16S rRNA gene abundance is *Listeria monocytogenes* – 14.1%, *Pseudomonas aeruginosa* – 4.2%, *Bacillus subtilis* – 17.4%, *Escherichia coli* – 10.1%, *Salmonella enterica* – 10.4%, *Lactobacillus fermentum* – 18.4%, *Enterococcus faecalis* – 9.9%, and *Staphylococcus aureus* – 15.5%.

The PowerPlant DNA extraction method was selected to verify its e□ciency at extracting accurate representative quantities of DNA from both Gram-positive and Gram-negative bacteria present in the mock microbial community standard (MC). Mock sap samples were prepared using 3-ml of bacterial-free xylem sap (previously filtered through a 0.22-μm filter and showing no amplification of 16S rRNA gene with PCR1, see below) to which we added 50 μl of ZymoBIOMICS Microbial Community Standard Cells (Zymo Research Corp., Irvine, CA, USA). Mock sap samples were then processed as indicated above for the PowerPlant DNA extraction protocol. After DNA extraction, the four PCR protocols described below were evaluated. There were four replicates per combination of PCR protocol.

### Effect of primer pairs to assess xylem bacterial composition of two olive cultivars

Olive groves from an experimental plot located at the Institute for Sustainable Agriculture from Spanish National Research Council (IAS-CSIC) facilities in Córdoba (southern Spain) were used to test the differences in xylem bacterial composition between two of the most widely grown olive cultivars in Spain. The orchard was established in September of 2014, with two-year old olive trees of cultivar ‘Picual’ and ‘Arbequina’ propagated at ‘Plantas Continental S.A.’ nursery (Ribero de Posadas, Córdoba, Spain). Both genotypes were planted in the orchard with a randomly block design and received similar growing practices until sampling.

Seven trees of each cultivar were sampled in May 2017, and xylem sap was extracted as described above. The average sap volume extracted ranged from 3.5 to 4 ml per sampled branch. In this case, and to facilitate DNA extraction, the sap samples were immediately filtered through a 0.22-μm Millipore filter, and the filters retaining all microbial cells contained in the sap and the filtered sap were stored independently at −20°C or −80°C, respectively. DNA extraction was performed using the PowerPlant protocol. First, the filtered cells were resuspended in the extraction buffer by vortexing briefly and then the microbial suspension was processed following manufacturer’s instructions (**Table 1**). To analyze bacterial communities, DNA samples were amplified using the four PCR protocols described below.

### 16S rRNA gene amplification

Four PCR protocols using five primer pairs targeting different hypervariable regions of the 16S rRNA gene were compared to evaluate their performance in metabarcoding studies of xylem sap bacterial communities. Those primer pairs have been described in previous studies as appropriate to avoid co-amplification of plant chloroplast and mitochondrial DNA. For that, two primer pairs we were used in two direct PCR protocols and additionally three primer pairs were used in two nested PCR protocols to evaluate if an increase in amplification yield could be obtained by the nested-PCR approach (**Table 2**):

**Table 2.**
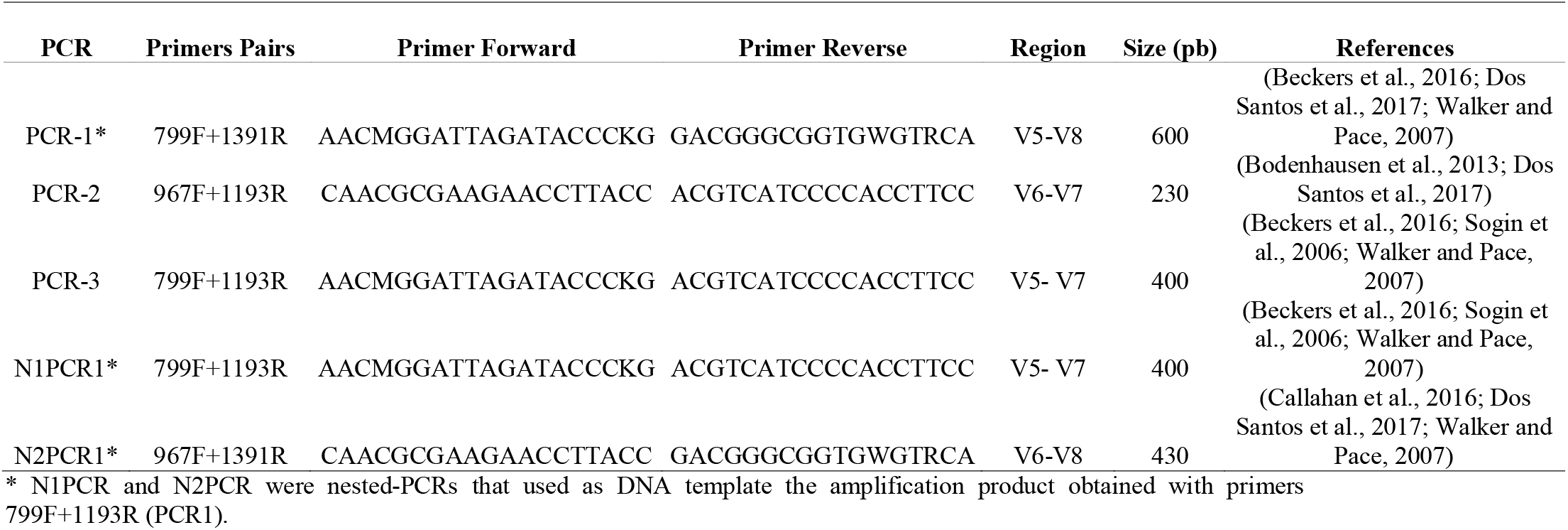
PCR protocols used in the study, with primer sequences, hypervariable region of 16S rRNA gene amplified and expected product size.

PCR1-(799F/1391R) uses primers 799F and 1391R that amplify a 600 bp fragment of the V5-V8 region of the bacterial 16S rRNA gene (Chelius and Triplett, 2001; Walker and Pace, 2007; Beckers et al., 2016) and its amplification product is used as a template for the second round of nested-PCRs described below.

PCR2-(967F/1193R) uses primers 967F and 1193R that amplify the V6-V7 region of the bacterial 16S gene (Dos Santos et al., 2017). This PCR was initially selected to test the different DNA extraction protocols, and it was also included to compare its performance with the other three PCR protocols described below.

PCR3-(799F/1193R) uses primers 799F and 1193R that amplify the V5-V7 region of the bacterial 16S rRNA gene (Chelius and Triplett, 2001; Bodenhausen et al., 2013; Beckers et al., 2016).

N1PCR1-(799F/1391R+799F/1193R) uses primers 799F and 1391R in the first round of PCR (i.e., PCR1), and primers 799F and 1193R in the second round of PCR.

N2PCR1-(799F/1391R+967F/1391R) uses primers 799F and 1391R in the first round of (i.e., PCR1), and primers 967F and 1391R that amplify the V6-V8 region of the bacterial 16S rRNA gene in the second round of PCR (Sogin et al., 2006; Walker and Pace, 2007; Beckers et al., 2016).

All PCRs were carried out in 25 μl reaction volumes containing 0.05 units of MyTaq™ DNA Polymerase (Bioline Laboratories, London, UK), 1x MyTaq™ Mix (Bioline) and forward and reverse primers at a concentration of 0.3 μM each, and 3μl of template DNA. PCR protocol consisted of an initial denaturalization step at 95°C for 5 min, followed by 35 cycles of 95°C for 1 min, 53°C for 45 sec, 72°C for 1 min, and a final elongation step at 72°C for 8 min. For the nested-PCR approach, 1 ul of the amplification product of the first reaction was used as template for the second PCR that was performed with same conditions as those of the first PCR.

### Library preparation and sequencing

PCR amplicons were cleaned up before adaptor addition using the Ampure XP magnetic bead system (Beckmann-Coulter, MA, USA) according to manufacturer’s recommendation. Dual barcode indices and sequencing adaptors were attached to each amplicon using the Illumina Nextera XT Index kit (Illumina, Inc., San Diego, CA, USA) following manufacturer’s protocol, followed by a further Ampure XP cleanup step. Purified amplicons were quantified using the Quant-iT™ PicoGreen™ dsDNA Assay Kit (Thermo Fisher Scientific) and a Tecan Safire microplate reader (Tecan Group, Männedorf, Switzerland). Equimolecular amounts from each individual sample in 10 mM of Tris were combined and the pooled library was additionally purified with two rounds of Ampure XP cleanup step. The library was sequenced by the Genomics Unit at ‘Fundación Parque Científico de Madrid’ (Madrid, Spain) using the Illumina MiSeq platform (Nano-V2; PE 2x 250 bp). The ZymoBIOMICS microbial standard (Zymo Research Corp., Irvine, CA, USA) and water (no template DNA) were used as internal positive and negative controls, respectively, for library construction and sequencing. Raw sequence data have been deposited in the Sequence Read Archive (SRA) database at the NCBI under BioProject accession number PRJNA684121.

### Data processing and bioinformatics analysis

16S rRNA gene sequences were analyzed and classified using the Quantitative Insights into Microbial Ecology bioinformatics pipeline, QIIME2 (version 2019.10; https://view.qiime2.org/; Caporaso et al., 2010; Bolyen et al., 2018) with default parameters unless otherwise noted. Demultiplexed sequences were imported as CASAVA format. Sequence quality control, denoising and chimeric filtering was performed with DADA2 pipeline (Callahan et al., 2016). Taxonomy affiliation was identified by operational taxonomic units (OTUs) at 99% similarity using VSEARCH consensus taxonomy classifier (Rognes et al., 2016) based on Greengenes_13_8_99 (DeSantis et al., 2006; McDonald et al., 2011) and Silva_132_99 (Quast et al., 2013; Yilmaz et al., 2014) reference databases. Singletons were discarded for downstream analysis.

Alpha and beta diversity as well as alpha rarefaction curves were conducted rarefying all samples to the minimum number of reads found. Sequencing depth was of 2,276 and 2,489 for the evaluation of the DNA extraction protocols and the microbiome of the olive tree cultivars, respectively. Rarefaction curves and alpha-diversity indexes (Shannon, and Richness or number of observed OTUs) were performed using the OTU frequency matrixes at the OTU level with the online tool MicrobiomeAnalyst (https://www.microbiomeanalyst.ca; Chong et al., 2020). The Kruskal-Wallis test (*P* <0.05) with FDR correction (Benjamini and Hochberg, 1995) was used to determine the effects of the factors in the study on alpha diversity among the studied factors. To analyze the association of PCR protocols and olive genotypes with these alpha diversity matrices, we applied General Linear Modelling (GLM) using the lme4 package in R (Bates et al., 2015), with a two factors factorial design, being the PCR protocols [967F-1193R (PCR2), n=14; 799F-1193R (PCR3), n=14; 799F-1391R + 799F-1193R (N1PCR1), n=14; 799F-1391R + 967F-1391R (N2PCR1), n=14], and the olive genotype (‘Arbequina’; n= 28; ‘Picual’, n= 28) the two main factors. Venn diagrams were generated using the “Venn diagram” online tool (http://bioinformatics.psb.ugent.be/webtools/Venn/) and were used to identify shared (core microbiome) or unique taxa according to the DNA extraction protocol or the PCR primers used.

For beta diversity analysis, we performed multivariate hierarchical clustering analysis (using Bray-Curtis index to measure distance and the Ward clustering algorithm), as well as non-supervised principal component analysis (PCoA) using Bray-Curtis distance (a non-phylogenetic metric) (Beals, 1984), and the weighted UniFrac distances (a phylogenetic metric) (Lozupone and Knight, 2005) were performed using the OTU frequency matrixes at the OTU level with QIIME2 to test for similarities among the bacterial communities according to the DNA extraction protocol, or among the PCR protocols and between the olive genotypes. In addition, the PERMANOVA test (*P* <0.05) was used to determine effects of those factors.

The theoretical relative abundance of the ZymoBIOMICS Microbial Community Standard was compared with the estimated relative abundances of the identified operational taxonomic units (OTUs) at genus level that were obtained for each PCR and reference database (Silva or Greengenes) using Spearman’s correlation analysis (McGovern et al., 2018). Similarity was considered as significant if *P* value < 0.05 and as a trend if 0.05 ≥ *P* value ≤ 0.1.

Finally, to analyze in further detail the differences in microbiome composition at genus level among the different PCR protocols, a negative binomial model approach based in the DESeq2 package in R (Love et al., 2014) was used. Wald tests were performed and only genera remaining significant (*P*<0.01) were retained.

## Results

### Effect of DNA extraction protocols on xylem sap bacterial community assessment

In the bacterial community analyses of the 12 different DNA extraction protocols tested a total of 108,236 high-quality 16S rRNA gene paired-end sequences with an average of 4,510 sequences per DNA protocol were retained after discarding poor quality sequences. From those, approximately 56% could be classified into bacterial operational taxonomic units (OTUs) with a mean of 2,460 and 2,468 bacterial sequences in Greengenes_13.8 or Silva_132 databases, respectively (**Table 3**). However, the number of plant organellar rRNA gene sequences amplified varied according to the DNA extraction kit, with the NorgenMicrobiomeV2 and PureLink DNA extraction kits showing the highest proportion of organellar reads (≥70%) and both NucleoSpin and the PowerLyzer DNA extraction kits showing the lowest proportion (≤4%), independently of the reference database used for taxonomic assignation (**Table 3**). A total of 209 or 248 OTUs, with an average number of 58 or 60 OTUs, were identified as bacteria when using the Greengenes_13.8 or Silva_132 databases, respectively (**Table 3**), with the higher number of OTUs being identified when using the NucleoSpin and PowerLyzer DNA extraction kits (i.e., identified OTUs raged between 84 to 93) (**Table 3**).

**Table 3.**
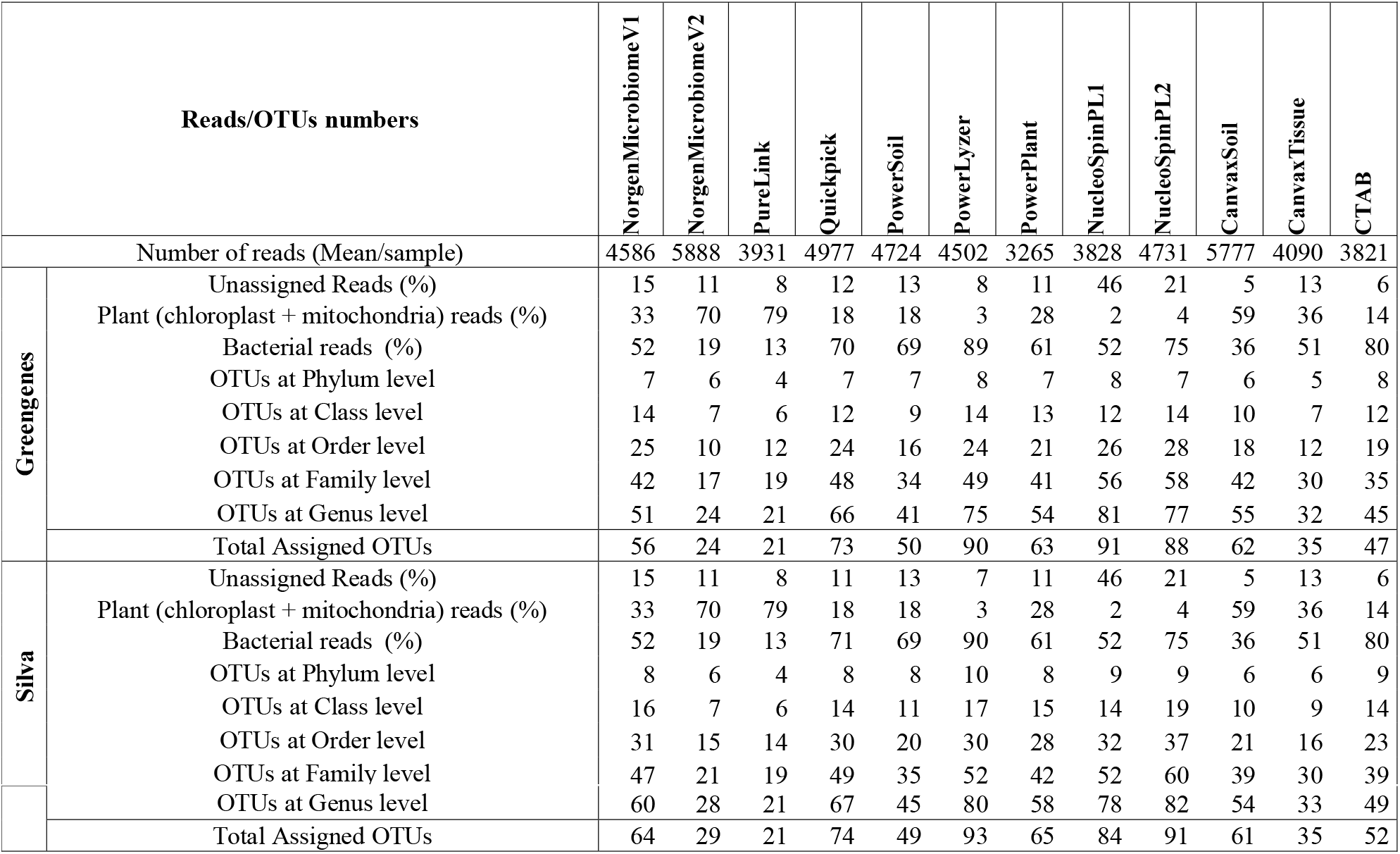
Number of reads and OTUs derived from NGS analysis of xylem sap microbiome from DNA samples extracted with 12 extraction protocols, amplified with the 967F+1193R primer pair (PCR2) and taxonomic assignation using the Greengenes 13-8 and Silva 132 reference databases

Rarefaction curves of observed OTUs (Richness) indicated a good sequencing coverage among all DNA extraction protocols with no differences between both reference databases (**Suppl. Fig. 1**). DNA extraction protocol significantly affected xylem sap bacterial community assessment. Thus, Richness alpha diversity index showed significant differences (*P*<0.044) among the different DNA extraction methods, whereas no significant differences (*P*>0.091) were found for the Shannon index, irrespective of the taxonomy database used (**Figure 1**). On the other hand, hierarchical clustering analysis and PCoA of Bray-curtis index and weighted UNIFRAC distances using OTU frequencies differentiated xylem bacterial communities in four clusters according to the DNA extraction method, irrespective of the database used for taxonomy assignation (**Figure 2A**). Thus, most DNA extraction kits from four different brands clustered together (Group 4), whereas the CTAB protocol (Group 1), both Canvax kits (Group 2), and the two NorgenMicrobiome protocols (Group 3), clustered independently from each other; although the Group 1 and Group 2 clustered closer and apart from the other DNA extraction kits (**Figure 2A**). Similarly, PCoA of Bray-Curtis and weighted UNIFRAC distances differentiated xylem bacterial communities according to the DNA extraction kits in four groups (**Figure 2B**), irrespective of the database used. PERMANOVA indicated a significant clustering due to the DNA extraction protocol used (*pseudo-F* < 12.386; *P* = 0.001). This clustering, derived from beta-diversity analyses, of the 12 DNA extraction kits in four independent groups was used to summarize the influence of DNA extraction kits in assessing the composition and abundance of bacterial communities on olive xylem sap.

**Figure 1.**
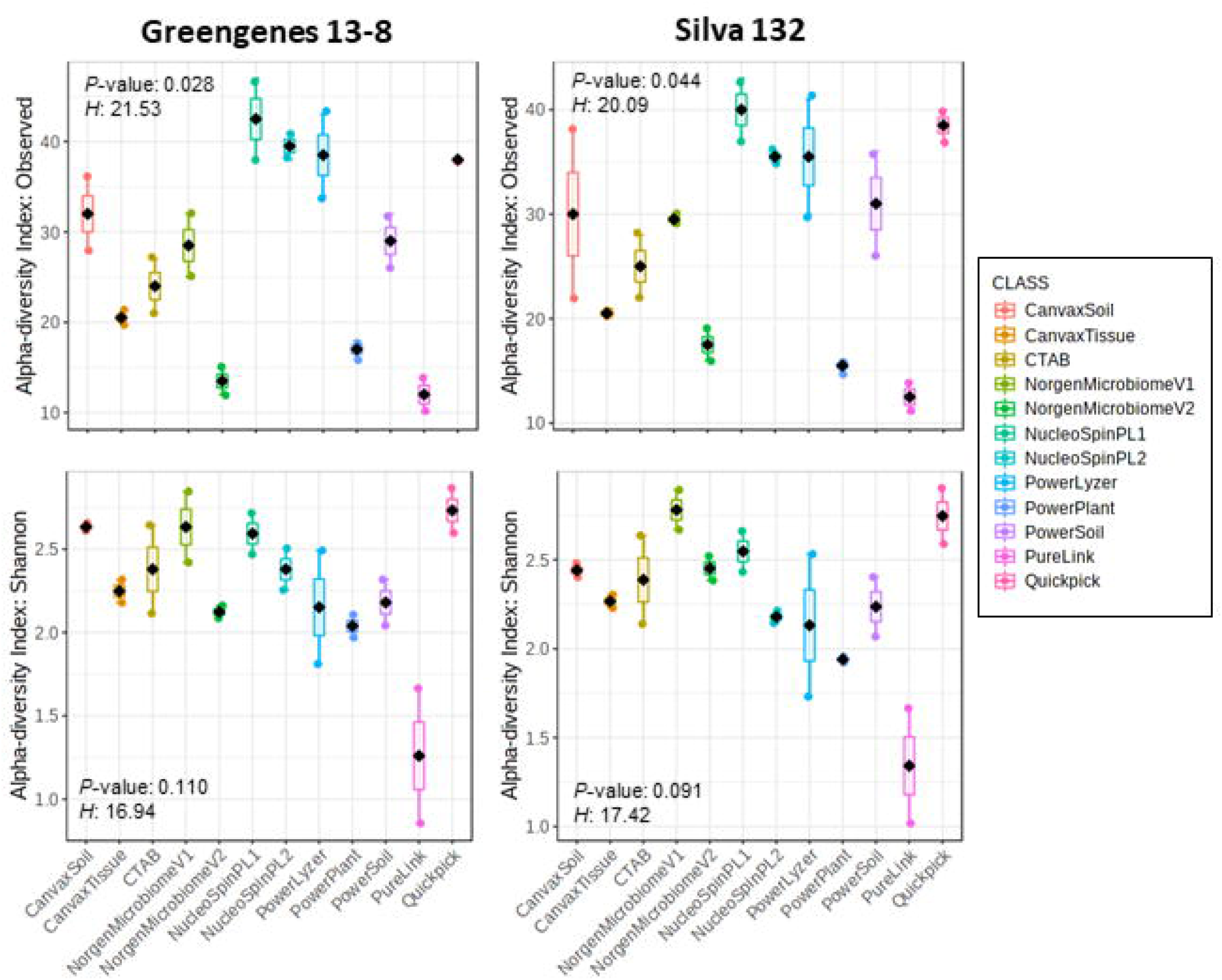
Boxplots of Richness and Shannon alpha-diversity indices of olive xylem bacterial communities at OTU taxonomic level determined by different DNA extraction kits and after taxonomic assignments with the Greengenes_13-8 and Silva_132 databases. Boxes represent the interquartile range while the black dots inside the box defines the median and whiskers represent the lowest and highest values. *P*-value was calculated using Kruskal-Wallis test at *P* < 0.05.

**Figure 2.**
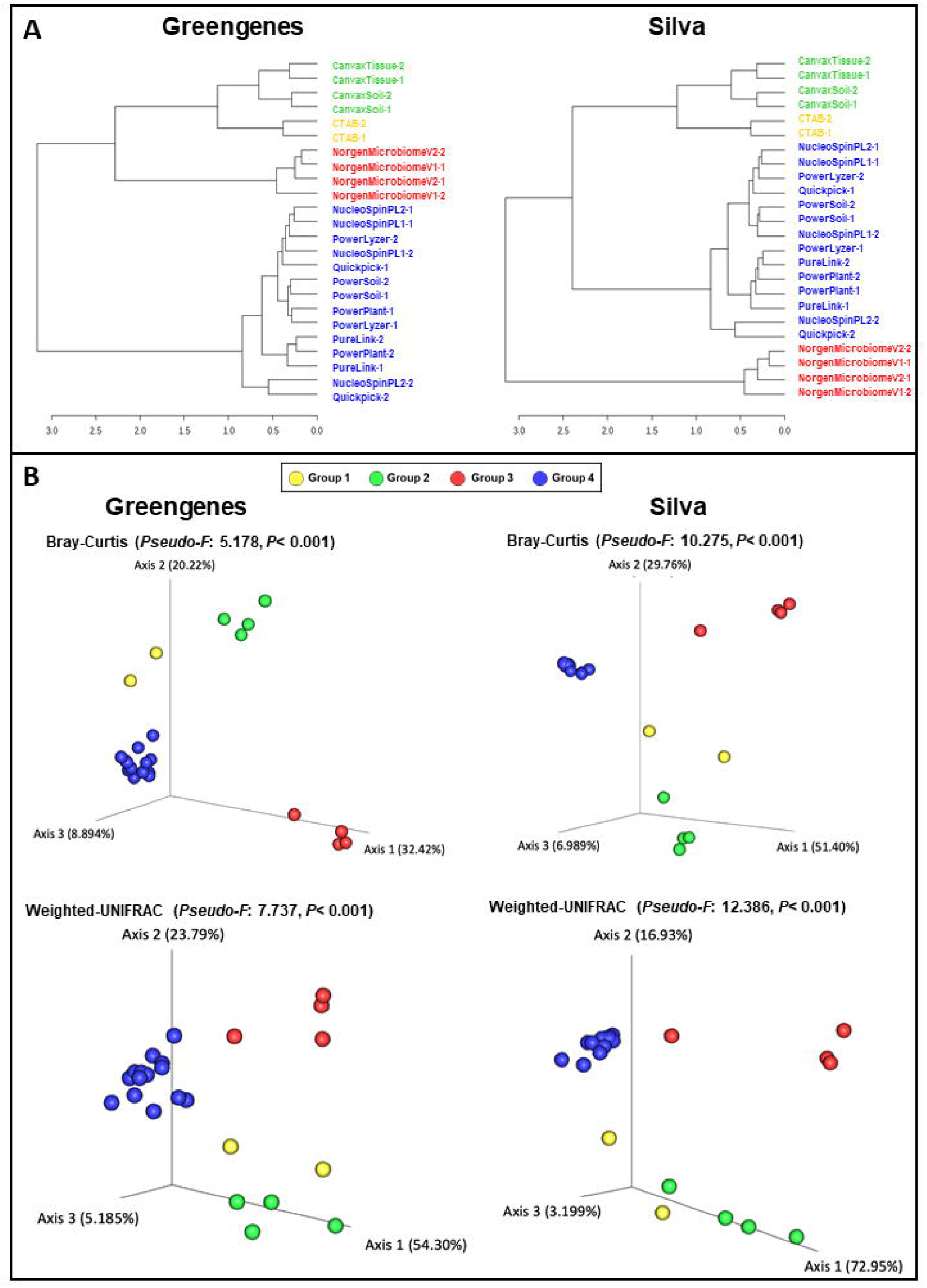
Hierarchical clustering dendrogram analysis using Ward method and Bray-Curtis distance (A) and principal coordinates analysis (PCoA) of weighted UniFrac and Bray-Curtis distances (B) of olive xylem bacterial communities obtained by using different DNA extraction kits and after taxonomic assignments with the Greengenes_13-8 and Silva_132 databases. Colored dots represent the four clusters obtained in the hierarchical clustering analysis. PERMANOVA (999 permutations; *P* < 0.05) was performed to test significant differences according to DNA extraction kits.

A total of 13 or 16 phyla, 29 or 34 classes, 53 or 65 orders, 118 or 124 families, 209 or 248 genera were identified when using the Greengenes or Silva database, respectively (**Suppl. Figure 2**). When comparing results from both databases, the core number of taxa detected by all the DNA extraction kits was of four phyla, seven classes, and 12 orders, whereas 16 or 21 families, and 12 or 14 genera were detected when using the Greengenes or Silva database, respectively. The lowest number of phyla and classes were identified when using the Canvax DNA extraction kits (Group 2), whereas the lowest number of orders, families, and genera were identified with the CTAB DNA extraction protocol (Group 1). In all cases, the highest numbers of total and unique phyla, orders, classes, families and genera were identified for the DNA kits included in Group 4. Finally, the DNA extraction kits included in Group 3 and 4 shared the highest number of bacterial taxa (**Suppl. Figure 2**).

Four phyla including Proteobacteria, Actinobacteria, Firmicutes, and Bacteroidetes were the most abundant and were identified by all DNA extraction kits, representing 51.8%, 26.2%, 9.7% and 9.6% of the total, respectively (**Figure 3; Suppl. Figure 2**). In addition, three minority phyla (<0.11% abundance), Nitrospirae, Fibrobacteres and Chloroflexi, were only identified when using the NorgenMicrobiome DNA extraction kits (Group 3), whereas other three phyla, Saccharibacteria, Gemmatimonadetes and Verrucomicrobia showed a low frequency (0.2%) and were only identified when using the different DNA extraction kits from Group 4 (**Suppl. Figure 2**). Seven bacterial classes were detected by all DNA extraction kits, with Actinobacteria, Alphaproteobacteria, Betaproteobacteria, Bacilli, and Gammaproteobacteria, being the most abundant, representing 26.1%, 22.6%, 19.7%, and 9.18% of the total, respectively (**Figure 3; Suppl. Figure 2**). At family level, 16 or 21 families, depending on the reference database used, comprised the core bacterial microbiome of all DNA extraction kits, being Propionibacteriaceae, Bradyrhizobiaceae, Comamonadaceae and Chitinophagaceae the most abundant (mean frequency ranged between 5.9 and 18%) (**Figure 3; Suppl. Figure 2**). Finally at genus level, 12 or 14 genera, depending of the reference database (**Suppl. Figure 2**), comprised the core bacterial microbiome, being *Propionibacterium*, and unidentified Comamonadaceae, *Sediminibacterium*, and unidentified Rhizobiales, and unidentified Methylophilaceae, an unidentified Bradyrhizobiaceae, *Novosphingobium, Staphylococcus, Bradyrhizobium*, and *Flavobacterium* the most abundant genera (mean frequencies ranged from 18.8 to 3.0%) by all DNA extraction kits when using the Greengenes database; whereas with the Silva database *Propionibacterium, Bradyrhizobium*, an unidentified Comamonadaceae, an unidentified Chitinophagaceae, an unidentified Methylophilaceae, *Novosphingobium, Bosea, Flavobacterium, Staphylococcus* and *Pseudomonas* were the most abundant genera (mean frequencies ranged from 18.7 to 2.8%). Interestingly, up to 114 genera could be detected uniquely by the DNA extraction kits within Group 4.

**Figure 3.**
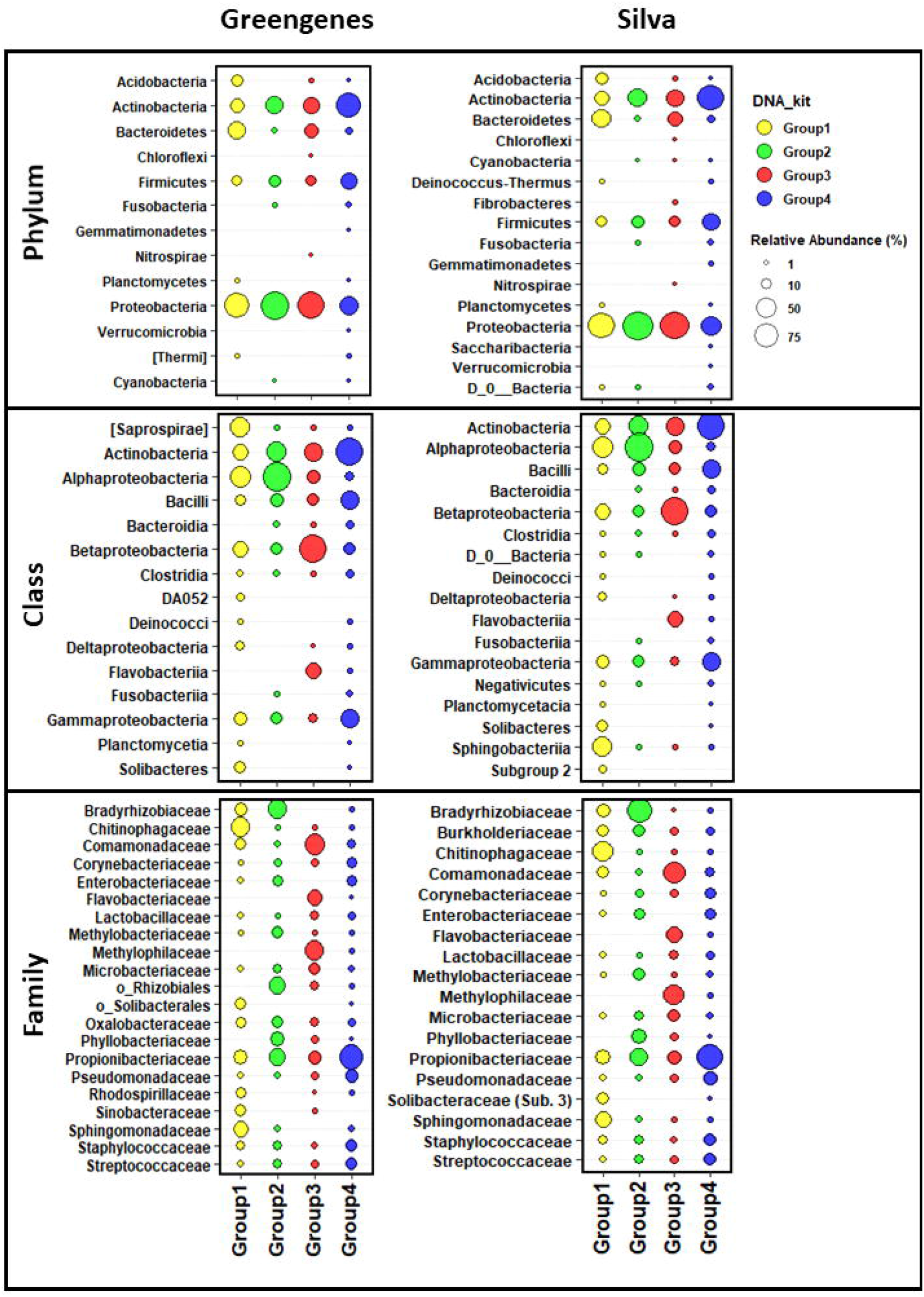
Taxonomic bubble plot of olive xylem bacteria relative abundance at phylum, class and family level present in four groups of DNA extraction kits and after taxonomic assignments with the Greengenes_13-8 and Silva_132 databases. Circle sizes represent the relative abundance and colors indicate DNA extraction kits groups shown in Figure 2. Only abundances greater than 1% are shown.

### Validation of a DNA extraction method, primer pairs performance and taxonomy reference databases

In this study, we used a mock microbial community standard to determine if a selected DNA extraction kit (the PowerPlant from Qiagen) within Group 4 (the one providing the highest values of alpha diversity) provides an accurate representation of the identified microbial communities and if this can be affected by the PCR protocol and the reference database used for taxonomic assignation (Greengenes 13-8 and Silva 132).

We found a very small background amplification of other OTUs or genera other than the eight expected to be present in the ZYMO mock community. Background amplification was observed on all samples and PCRs when Silva database was used and represented about 9.8 to 32.6% of reads. Among the background bacteria amplified unidentified OTUs belonging to the Class Bacilli, the Order Lactobacillales and the genus *Granulicatella* were detected, although most of ‘contaminant’ reads were assigned only as Bacteria and could not be assigned taxonomically to any phyla. On the other hand, when using Greengenes database, background amplification was detected only on PCR2 and N1PCR1 but represented less than 2% of reads with only the genus *Granulicatella* being detected.

All eight expected bacterial genera were detected by the four PCR protocols used irrespective of the reference database used (**Figure 4**). However, there was a large effect on accuracy of genera relative abundances estimation depending on the PCR and the reference database used. Thus, only when using primer pair 799F/1193R (PCR3) and when performing the taxonomic assignation with the Silva 132 database, a significant correlation (Spearman coefficient 0.714, *P*=0.046) was found between the theoretical and the estimated bacterial community composition (**Figure 4**). Consequently, we decided to select the Silva database for taxonomic assignation for further analysis.

**Figure 4.**
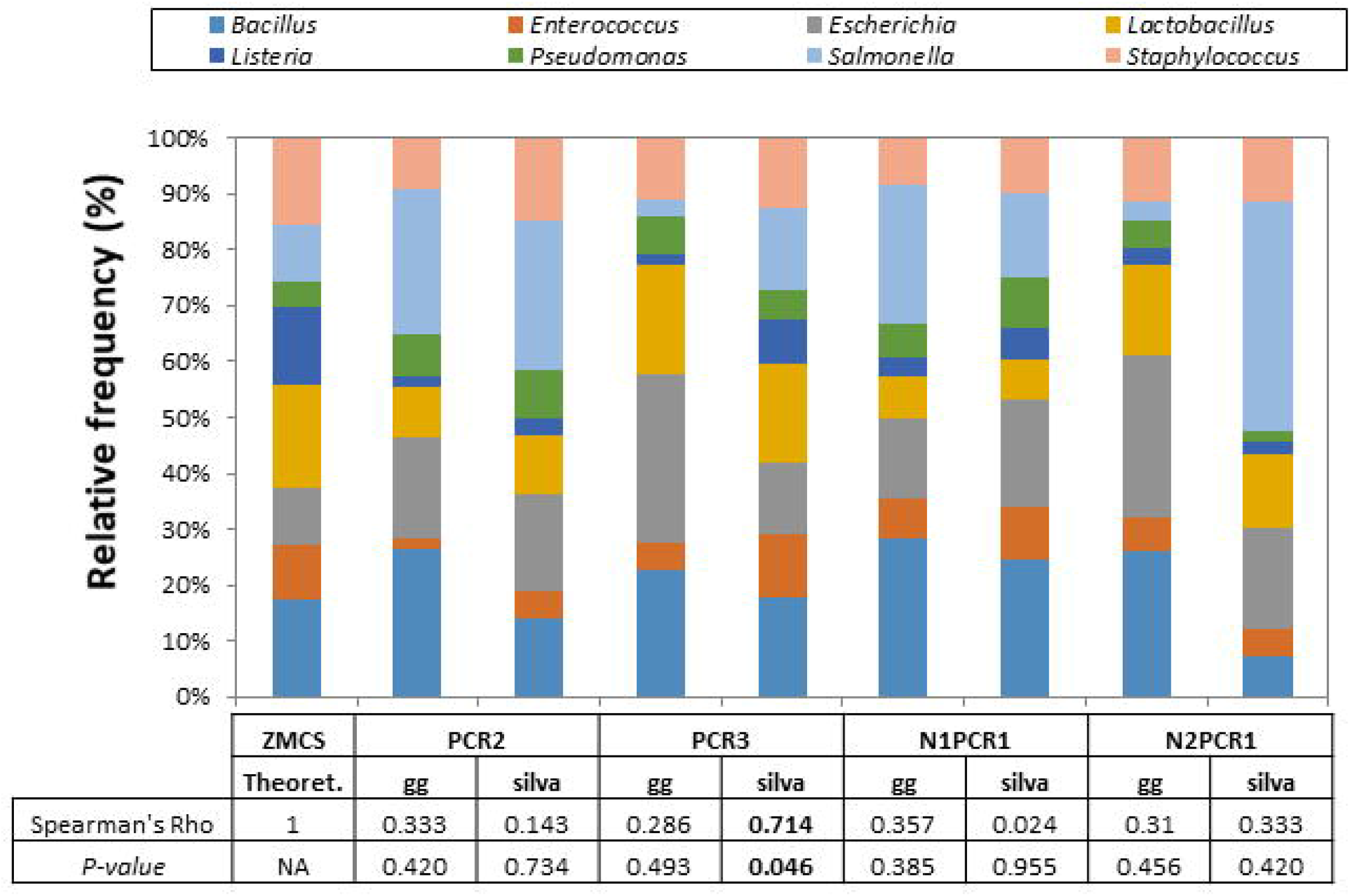
Relative abundance of ZymoBIOMICS Microbial Community Standard (ZMCS) composition according to the PCR primer pairs tested and the database used for taxonomic assignment [Greengenes_13-8 (gg) and Silva_132 (silva)]. Comparison with the theoretical abundances of ZMCS was based on spearman correlation coefficient.

### Effect of primer pairs on xylem sap bacterial community assessment of two olive cultivars

For the bacterial community analyses of the 14 samples from ‘Picual’ and ‘Arbequina’ olive plants, after screening our data for poor quality sequences and removing chimeras and unassigned reads, we recovered a total of 86,489, 54,143, 44,444 and 33,261 high-quality paired-end sequences with the PCR2, PCR3, N1PCR1 and N2PCR1 protocols, respectively; with an average of 6,178, 3,867, 3,175 and 2,376 sequences per PCR, respectively (**Table 4**). The number of unassigned reads was lower (<3%) for the two direct PCRs (PCR2 and PCR3) as compared to the two nested PCRs (>6.5%). However, and only for primers 967F-1193R (PCR2) a high percentage of reads were of plant origin, whereas no such amplification occurred with the other PCR protocols (**Table 4**).

**Table 4.**
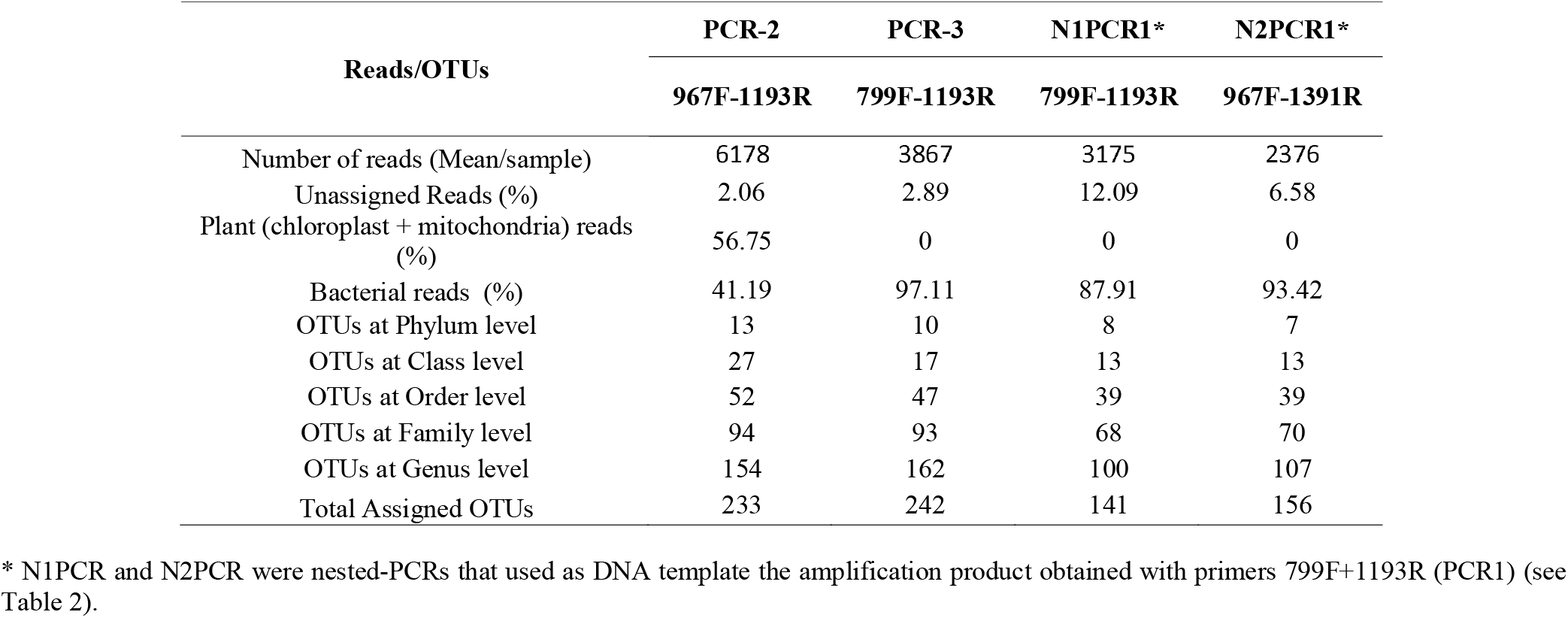
Number of reads and OTUs derived from NGS analysis of xylem sap microbiome from DNA samples extracted with the PowerPlant protocol, amplified with four PCR protocols and taxonomic assignation using the Silva 132 reference database.

| Reads/OTUs | PCR-2 | PCR-3 | N1PCR1* | N2PCR1* |
| --- | --- | --- | --- | --- |
|  | 967F-1193R | 799F-1193R | 799F-1193R | 967F-1391R |
| Number of reads (Mean/sample) | 6178 | 3867 | 3175 | 2376 |
| Unassigned Reads (%) | 2.06 | 2.89 | 12.09 | 6.58 |
| Plant (chloroplast + mitochondria) reads (%) | 56.75 | 0 | 0 | 0 |
| Bacterial reads (%) | 41.19 | 97.11 | 87.91 | 93.42 |
| OTUs at Phylum level | 13 | 10 | 8 | 7 |
| OTUs at Class level | 27 | 17 | 13 | 13 |
| OTUs at Order level | 52 | 47 | 39 | 39 |
| OTUs at Family level | 94 | 93 | 68 | 70 |
| OTUs at Genus level | 154 | 162 | 100 | 107 |
| Total Assigned OTUs | 233 | 242 | 141 | 156 |

\* N1PCR and N2PCR were nested-PCRs that used as DNA template the amplification product obtained with primers 799F+1193R (PCR1) (see Table 2).

For Richness alpha diversity index, we found a significant effect for the PCR protocol (*P*<0.001) and no effect for olive genotype (*P*=0.537). For Shannon diversity index, no significant effect was found for any of the factors (*P*>0.554) (**Figure 5**). In general, both nested-PCR approaches showed lower values of alpha diversity indexes (**Figure 5**). To determine whether choice of PCR approach (primer pair) influenced microbial community composition, we calculated two beta diversity metrics (Bray-Curtis and weighted UNIFRAC) and included the PCR protocol, olive genotype and the interaction PCR protocol*olive genotype as explanatory variables in PERMANOVA models. In these analyses, microbial beta diversity estimates were significantly influenced by the PCR protocol (Bray-Curtis: pseudo-*F*= 22.12, *P*<0.001; weighted-UNIFRAC: pseudo-*F*= 11.96, *P*<0.001), as well by the interaction PCR protocol*genotype (Bray-Curtis: pseudo-*F*= 10.22, *P*<0.001; weighted-UNIFRAC: pseudo-*F*=5.93, *P*<0.001), but not by the olive genotype (Bray-Curtis: pseudo-*F*= 0.53, *P*=0.90; weighted-UNIFRAC: pseudo-*F*=1.61, *P*=0.118) on microbial beta diversity estimates (**Figure 6**). N1PCR1, that is a nested-PCR that uses in the second round of PCR same primers than those used in PCR3, clustered and grouped together and closer to samples amplified with this later PCR both in the hierarchical cluster analysis (**Figure 6A**) and in the PCoA analysis using Bray-Curtis distance (**Figure 6B**). According to the phylogenetic distances among the bacterial communities amplified by each PCR protocol, primers 967F-1193R (PCR2) showed a more distinct bacterial community composition as compared to PCR3, N1PCR1 and N2PCR1 that tended to overlap (**Figure 6B**).

**Figure 5.**
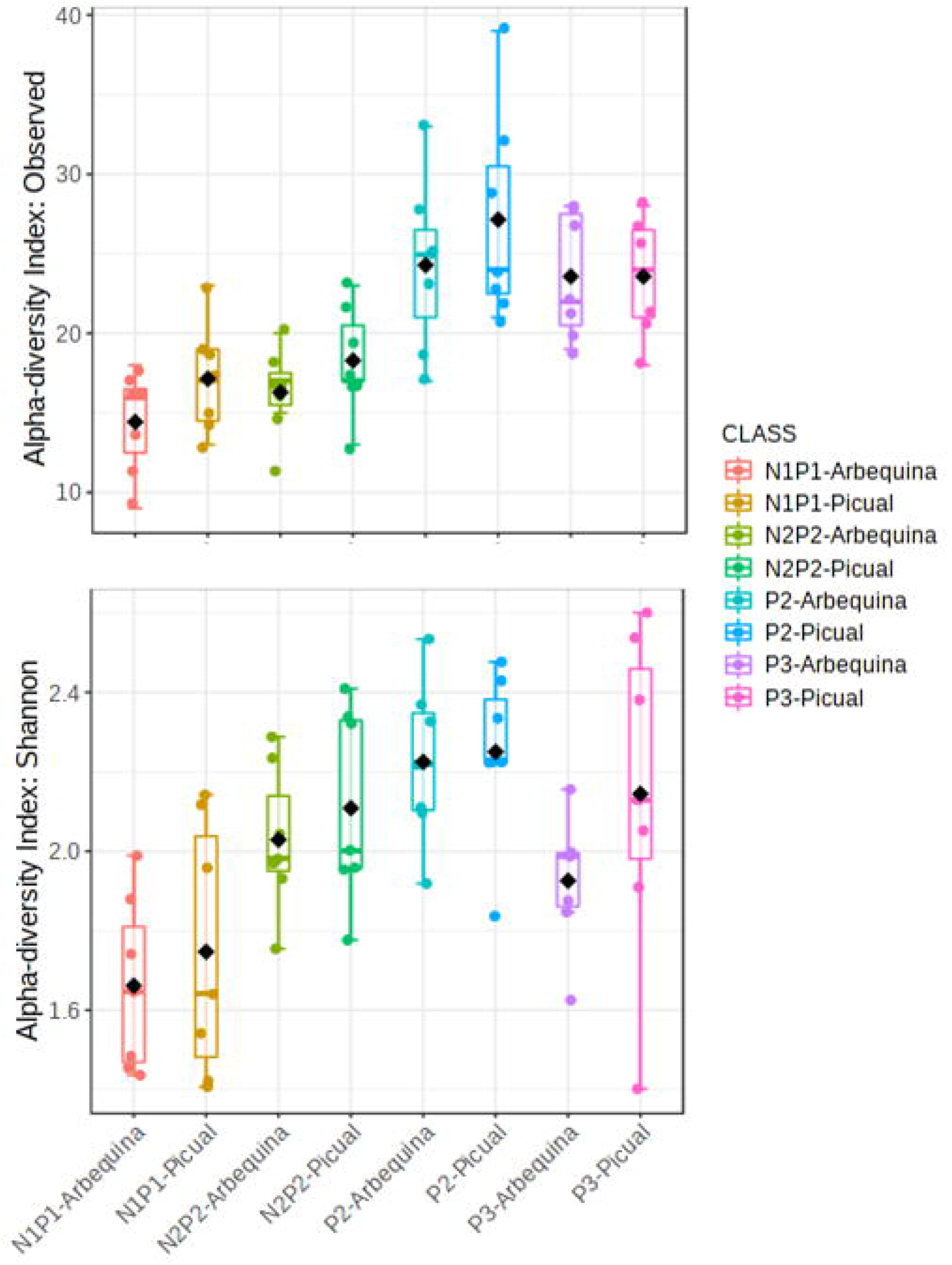
Boxplots of Richness and Shannon alpha-diversity indices of xylem bacterial communities from ‘Picual’ and ‘Arbequina’ olive cultivars at OTU taxonomic level determined by using different PCR primer pairs. Boxes represent the interquartile range while the black dots inside the box defines the median and whiskers represent the lowest and highest values. *P*-value was calculated using General Linear Modelling (GLM).

**Figure 6.**
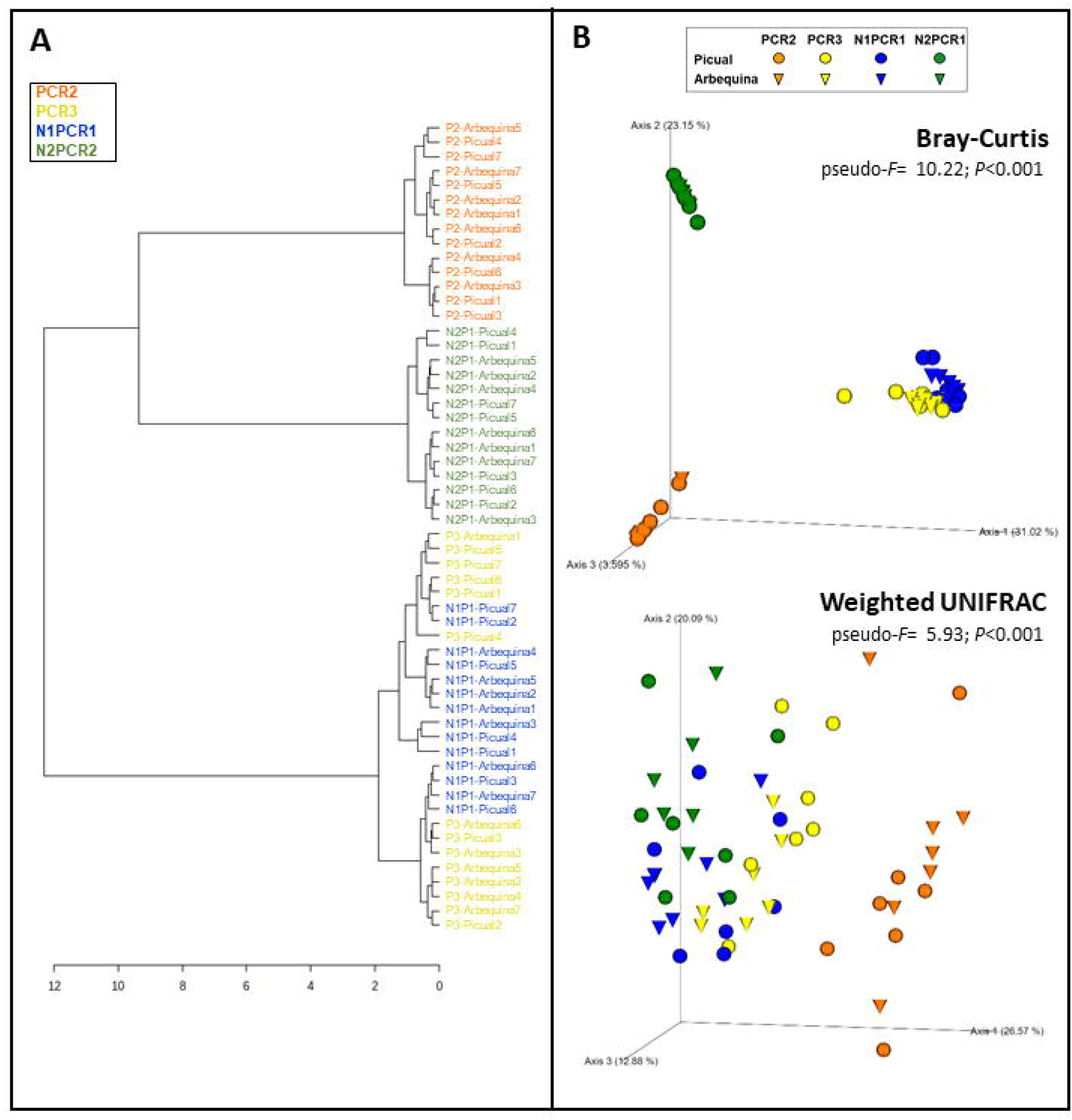
Hierarchical clustering dendrogram analysis using Ward method and Bray-Curtis distance (A) and principal coordinates analysis (PCoA) of weighted UniFrac and Bray-Curtis distances (B) of xylem bacterial communities from ‘Picual’ and ‘Arbequina’ olive cultivars obtained by using different PCR primers pairs. PERMANOVA (999 permutations; *P* < 0.05) was performed to test significant differences according to PCR primers used and olive genotypes.

The differences on alpha and beta diversity indexes found among PCR protocols were due to differences on both the number of bacterial taxa identified and their abundances. In fact, a total of eight to 13 phyla, 13 to 27 classes, 39 to 52 orders, 68 to 94 families, 100 to 162 genera and 141 to 242 species were identified depending of the PCR protocol (**Table 4, Suppl. Figure 3**). The core number of taxa detected by all the PCR protocols included six phyla, nine classes, 28 orders, 48 families and 50 genera. The lowest number of all bacterial taxa were obtained for both nested-PCR protocols, whereas the highest numbers of phyla, classes, and orders was detected with primers 967F-1193R (PCR2), and those of genera and species was obtained with primers 799F-1193R (PCR3). In all cases, the highest numbers of unique and shared taxa occurred for both direct PCR protocols (PCR2 and PCR3) (**Table 4; Suppl. Figure 3**).

In general, the bacterial taxa with higher abundance were detected by all the four PCR protocols (**Figure 7; Suppl. Figure 3**). At phylum level Actinobacteria, Proteobacteria, Firmicutes, and Deinococcus-Thermus represented 51.9%, 29.0%, 16.8% and 1.2%, respectively. Six bacterial classes were detected by all PCR protocols, with Actinobacteria, Gammaproteobacteria, Bacilli, Alphaproteobacteria, Clostridia and Deinococci being the most abundant, representing 51.6%, 23.4%, 15.6%, and 5.5%, respectively (**Figure 7; Suppl. Figure 3**). Fourteen families were detected by all PCR protocols, with Propionibacteriaceae, Staphylococcaceae, Pseudomonadaceae, Burkholderiaceae, Enterobacteriaceae, Sphingomonadaceae, Corynebacteriaceae, Microbacteriaceae being the most abundant (mean frequency ranged between 42.4 and 3.8%, in that order) (**Figure 7; Suppl. Figure 3**).

**Figure 7.**
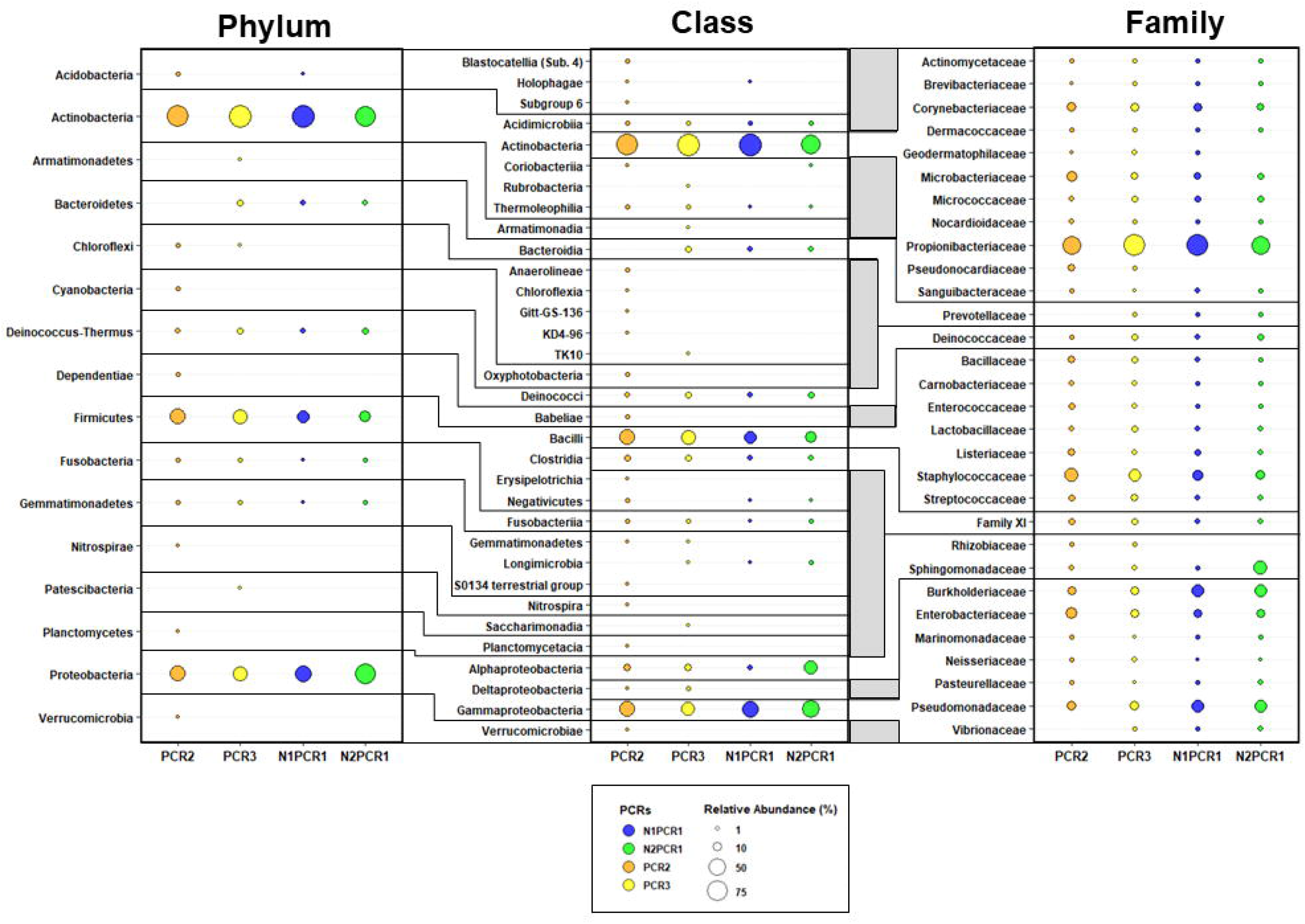
Taxonomic bubble plot of bacterial relative abundance at phylum, class and family level present in different PCR primers pair combination. Circle sizes represent the relative abundance and colors indicate PCR primers pair used. All phylum and class were shown while only the most abundant bacteria (98% of reads) at family level were represented. Horizontal lines indicate the taxonomy lineage from each bacterial family.

Finally, at genus level, 50 genera comprised the core bacteria, where the highest number of genera were detected by direct PCR primers (154 in PCR2 and 162 in PCR3) while the lowest number was obtained for both nested-PCR protocols (100 in N1PCR1 and 107 in N2PCR1). Direct PCR protocols showed the same unique number of genera (51 each one) while N2PCR1 displayed nine unique genera. No unique genera were found in N1PCR1 (**Table 4; Suppl. Figure 3**). Differential abundance analysis using DESeq2 displayed significant differences among genera between the each of the PCR primer pairs tested when compared to the selected PCR3 protocol. Thus, DESeq2 identified a significant and high enrichment (Log2 Fold Change > 5) of *Faecalibacterium, Prevotella, Geodermatophilus* and *Frigoribacterium* in PCR2, *Rhizobium, Enterobacter*, *Granulicatella* and *Brevibacterium* in N1PCR1 and *Brevundimonas*, *Staphylococcus, Prevotella* and *Dermacoccus* in N2PCR2 (**Figure 8**). Interestingly, *Enterobacter, Granulicatella, Prevotella* and *Brevibacterium* presented a significant higher abundance in all PCR protocols when comparing to PCR3 while *Pseudomonas* and *Pectobacterium* displayed an opposite behavior (**Figure 8**).

**Figure 8.**
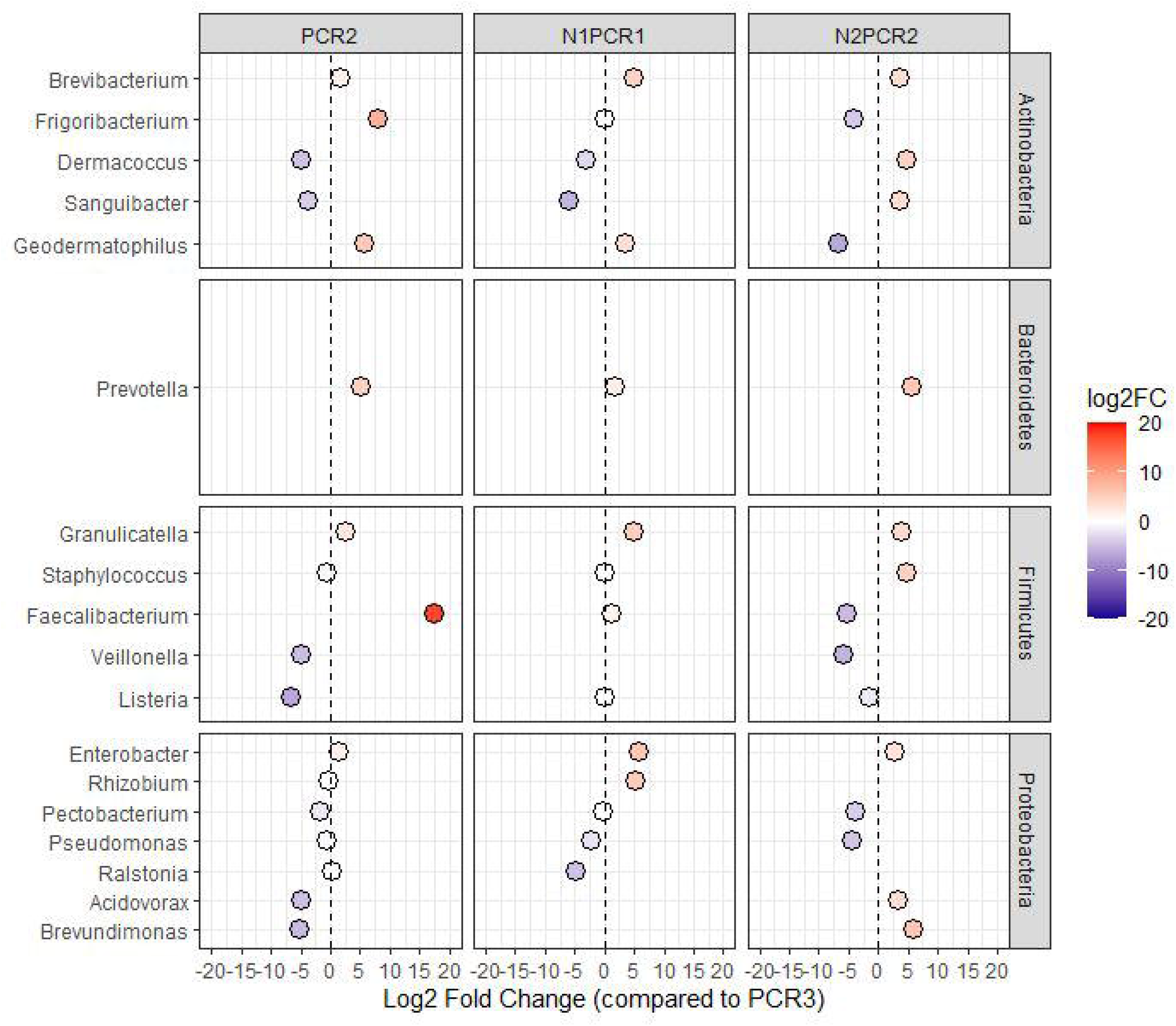
DESeq2 analysis of differentially enriched bacterial genera when using different PCR protocols. PCR2, N1PCR1 and N1PCR1 were compared against PCR3.The color scale bar indicates log2 fold change. Only significant genera (*P*<0.01) are shown.

## Discussion

Application of NGS to plant microbiome studies may lead to expand our understanding of the less easy to access and isolate microorganism, such as those present in the xylem vessels of plants. The exploration of xylem microbiome is essential to understand the diversity, biogeography and function of the host microbiota which may result in the development of innovative approaches based on microbiome exploitation that can contribute to protect plants against their pathogens (Knudsen et al., 2016). Microbiome research has increased considerably in recent years since sequencing is no longer the greatest impediment as it was in the past. However, the impact that the use of different commercially available DNA extraction kits and/or primers can exert in the assessment of the microbial diversity and community composition has received relative little attention to date (Kennedy et al., 2014).

Although assessment and optimization of different DNA extraction protocols should be one of the most important initial steps when developing a protocol for analysis of microbial communities in a new plant niche, due to its potential significant influence on the structure and diversity of the recovered community profile (Rubin et al., 2014), for all the studies assessing olive microbiome this assessment is lacking. Thus, although several studies have described olive endophytic microorganisms either after extraction from xylem woody chips (e.g., Keykhasaber et al., 2017; Anguita-Maeso et al., 2020; Giampetruzzi et al., 2020; Vergine et al., 2020), or from xylem sap when using a pressure chamber (Fausto et al., 2018; Sofo et al., 2019; Anguita-Maeso et al., 2020), none of those studies have examined the influence of DNA extraction kits or choice of primer pairs on microbiome characterization which might be essential to avoid possible bias on the results.

Nowadays, a wide range of commercial and ready-to-use DNA extraction kits are available from global life-sciences companies where cell lysis, washing and DNA capture are considered general steps among all of them. Aside from these similarities in the protocols, the commercial kits have some differences within these steps that included a distinct cell lysis procedure based on chemical lysis and/or mechanical cell disruption with bead beating, and on the DNA capture which is based either on a silica matrix in the presence of a high concentration of salt solution or the use of magnetic beads (Burbach et al., 2016). Additionally, the bead beating type (e.g., glass, garnet, ceramic, etc.) used in cell disruption must be considered in the DNA extraction procedure (Knudsen et al., 2016). In general, the vast majority of commercial DNA extraction kits include chemical and mechanical lysis methods together with silica membrane-based columns for DNA capture (e.g., DNeasy PowerPlant Pro Kit, DNeasy PowerLyzer PowerSoil Kit or PowerSoil^®^ DNA Isolation Kit).

After performing each DNA extraction protocol, we observed a noticeable differentiation between DNA yields obtained. Indeed, the quantity of DNA extracted with the commercial kit could be limited by the maximum DNA concentrations allowed by the filters used in the filtration step (Cruaud et al., 2014). From the 12 DNA kits assayed, we found PowerPlant as the most suitable DNA extraction kit for the characterization of olive xylem sap microbiome due to the integrity and concentration of DNA obtained, it is a short time-consuming procedure, the market price is within the lowest and the high percentage (92%) of sequences assigned to bacteria. These results are in line with other works that support its usefulness for subsequent microbiome analysis (Corcoll et al., 2017). However, PowerSoil kit also showed as good characteristics as those shown by the PowerPlant kit, although the time for extraction is a little bit longer. Consequently, it could also be considered useful for the analysis of olive microbiome, especially on studies in which other plant niches such as the rhizosphere may also be explored. In fact, this kit has gained special interest as the standard technique for extracting microbial DNA from environmental samples either soil or non-soil studies, including two of the largest microbiome initiatives, the Earth Microbiome Project and the Human Microbiome Project (Rubin et al., 2014).

Since bacterial 16S rRNA gene sequences present a high homology with chloroplast and mitochondrial rRNA genes (Sakai et al., 2004), the discrimination between host plant DNA, organellar rDNA and microbial 16S rDNA suppose an enormous challenge for the application of PCR-based methods in the study of plant microbiome (Beckers et al., 2016; Jackrel et al., 2017). Therefore, we determined the effect of primer pairs on xylem sap bacterial community characterization when using a mock- and an indigenous-microbial community extracted from two olive cultivars. The primers selected for our study had been reported to reduce co-amplification or organellar sequences being widely and commonly used for the analysis of plant-associated bacterial communities using Illumina MiSeq sequencing. The ZymoBIOMICS Microbial Community Standard, designed to assess bias and errors in DNA extraction protocols and to improve the quality and reproducibility of metagenomics analyses, was used to determine the efficacy of four primer pairs targeting different hypervariable regions of 16S rRNA gene for giving an accurate representation of the xylem microbial communities. This validation procedure has been used by other authors (e.g., McGovern et al., 2018; Neuberger-Castillo et al., 2020). The ZymoBIOMICS Microbial Community Standard offers a well-defined composition and an accurately characterized mock community consisting of three Gram-negative and five Gram-positive bacteria, easy and tough to lyse, respectively and two tough-to-lyse yeasts with varying sizes and cell wall composition. These wide range of organisms with different properties enables characterization, optimization, and validation of different lysis methods and PCR amplification protocols and it can guide construction of entire workflows being used as a routine quality control. This mock microbial DNA community standard allowed the identification of PCR3 (799F/1193R) as the most accurate primer pair in combination with the Silva 132 database for taxonomic assessment when comparing results with the theoretical composition of ZymoBIOMICS (**Figure 3**).

When using natural olive samples from ‘Arbequina’ and ‘Picual’ cultivars and comparing the four primer pairs, results indicated that primer pair 799F-1193R recovered the highest number of bacterial OTUs (242) and displayed a low co-amplification rate of organellar rRNA gene; although for several of the samples a removal of unspecific bands by agarose gel purification was needed prior to library sequencing. The Actinobacteria, Firmicutes and Proteobacteria phyla have been described as the most abundant in olive xylem sap by other authors (Sofo et al., 2019; Anguita-Maeso et al., 2020). In our study, although the same most abundant phyla, were detected by all PCRs; some phyla were detected exclusively by some PCR primers. For instance, PCR2 was the only one detecting Dependentiae, Verrucomicrobia, Nitrospirae, Planctomycetes and Cyanobacteria, whereas the Armatimonadetes and Patescibacteria phyla were exclusively detected by PCR3. At lower taxonomic level, 48 families formed the core microbiome among all PCR primer pairs tested. Among these, the families Propionibacteriaceae, Staphylococcaceae, Sphingomonadaceae, Burkholderiaceae, Enterobacteriaceae and Pseudomonadaceae were the most predominant although their relative abundance varied according to the PCR primer pairs used. Thus, PCR3 and N1PCR1 detected higher abundance of Propionibacteriaceae in comparison with PCR2 and N2PCR1. On the other hand, the families Staphylococcaceae and Enterobacteriaceae were detected at higher frequencies with PCR2 whereas the relative amount of Sphingomonadaceae was higher when using N2PCR1. Similarly, the families Burkholderiaceae and Pseudomonadaceae presented higher relative abundance when using nested-PCR primers instead of direct PCRs. Several of the core most abundant families found in our study such as Staphylococcaceae, Sphingomonadaceae, Burkholderiaceae, Enterobacteriaceae and Pseudomonadaceae have been detected in the xylem of different olive cultivars by other authors (Müller et al., 2015; Fausto et al., 2018; Sofo et al., 2019; Anguita-Maeso et al., 2020; Giampetruzzi et al., 2020).

The tendency to find low number of taxa in nested-PCRs when compared to direct PCR protocols was also observed at genus level. Although 50 genera composed the core bacterial microbiome, we found different unique genera depending on the PCR protocol. Interestingly, high number of unique genera was found in PCR2 and PCR3 (51 each one) whereas N2PCR1 showed nine exclusive genera and no one was found unique in N1PCR1. Differential abundance analysis showed distinct enrichment of some genera based on the PCR primer pairs used. In such way, when comparing PCR3 against the other three PCR protocols, we observed a significant enrichment of *Faecalibacterium, Prevotella, Geodermatophilus* and *Frigoribacterium* in PCR2; *Rhizobium, Enterobacter, Granulicatella* and *Brevibacterium* in N1PCR1 and *Brevundimonas, Staphylococcus, Prevotella and Dermacoccus* in N2PCR2. Within this comparison, *Faecalibacterium, Enterobacter* and *Brevundimonas* in PCR2, N1PCR1 and N2PCR2, respectively, displayed the greatest values of enrichment. These genera have been detected in other plant niches previously such as in rhizosphere or phyllosphere (Teixeira et al., 2010; Compant et al., 2019). Among them, *Brevundimonas* have been already described to confer fitness advantages to host plants due to its potential to act as soil bioremediator and plant growth promotor (Kumar and Gera, 2014; Singh et al., 2016) although its use as biological control agent against plant diseases is compromised due to the human pathogenic activity presented by some members of this genus (Ryan and Pembroke, 2018). On the other hand, we observed differences in ‘Picual’ and ‘Arbequina’ olive genotypes where the phyla *Acidobacteria* and *Gemmatimonadetes* were only present in ‘Picual’ cultivar. However, an in-depth study targeting a wide range of olive cultivars is needed to better understand the effect of olive genotype in shaping the xylem microbiome.

Our study demonstrates significant and noticeable differences among DNA extraction kits and PCR primers that influence the interpretation of the microbial community composition of olive xylem sap. Overall, our findings provide new insights and an integrated assessment of both the benefits and drawbacks of several commercially available DNA extraction kits and offer a guidance to other researchers in the choice of best-suited kits, taking into account cell lysis efficacy, DNA yield, microbial diversity recovered, processing time, and cost-effectiveness. Also, this study highlights the crucial choice of a good primer set to provide a nonbiased vision of the true composition of the analyzed microbial community avoiding the co-amplification of plant organellar rRNA genes. Our results, offered the roadmap to design an optimized strategy for selecting the most suitable PCR primer pair for assessing bacterial communities based on the use of artificial commercially-available mock community together with a precise and accurate bioinformatic workflow that can be followed when optimizing protocols for accurate depiction of the bacterial communities present in xylem vessels, or other plant niches.

## Acknowledgments

We acknowledge support of the publication fee by the CSIC Open Access Publication Support Initiative through its Unit of Information Resources for Research (URICI).

## Data Availability Statement

The raw sequence data have been deposited in the Sequence Read Archive (SRA) database at the NCBI under BioProject accession number PRJNA684121.

## Supplementary Material

**Figure S1**. Richness rarefaction curves at OTU taxonomic level obtained when using different DNA extraction kits and after taxonomic assignments with the Greengenes_13-8 and Silva_132 databases. Error bars show the standard deviation.

**Figure S2**. Prevalence Venn diagram showing the unique and shared bacterial taxa at phylum, class, order, family and genera level using the four clustered DNA extraction kits obtained shown in Figure 2. For each taxa, the venn diagram is shown using the Greengenes 13-8 and Silva_132 databases. Tables show the bacterial taxonomy interaction within each reference database.

**Figure S3**. Prevalence Venn diagram showing the unique and shared bacterial taxa obtained at phylum, class, order, family and genera level using the four PCR protocols. Tables show the bacterial taxonomy interaction within each PCR protocol.

