## Supplementary material for "Evaluation of established methods for DNA extraction and primer pairs targeting 16S rRNA gene for bacterial microbiome profiling of olive xylem sap": Suppl Figure S3

#### Phylum level

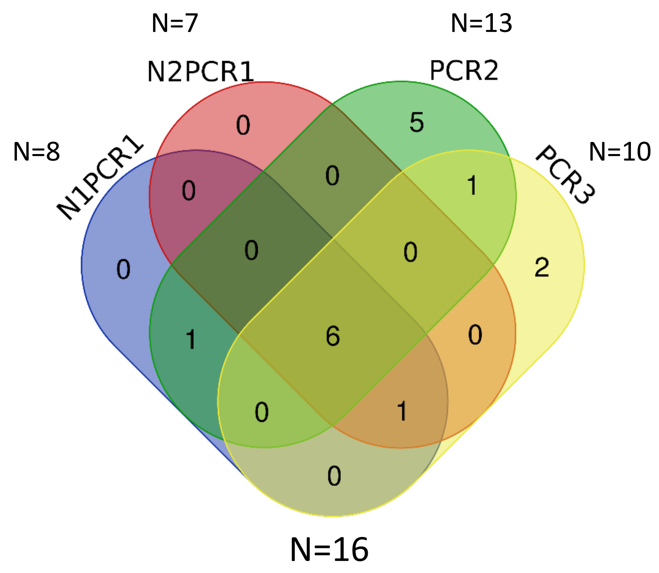


| PCRs | Total | Phylum |
|---|---|---|
| N1PCR1 N2PCR1 PCR2 PCR3 | 6 | Actinobacteria, Firmicutes, Deinococcus-Thermus, Proteobacteria, Fusobacteria, Gemmatimonadetes |
| N1PCR1 N2PCR1 PCR3 | 1 | Bacteroidetes |
| N1PCR1 PCR2 | 1 | Acidobacteria |
| PCR2 PCR3 | 1 | Chloroflexi |
| PCR2 | 5 | Dependentiae, Verrucomicrobia, Nitrospirae, Planctomycetes, Cyanobacteria |
| PCR3 | 2 | Armatimonadetes, Patescibacteria |

#### Class level

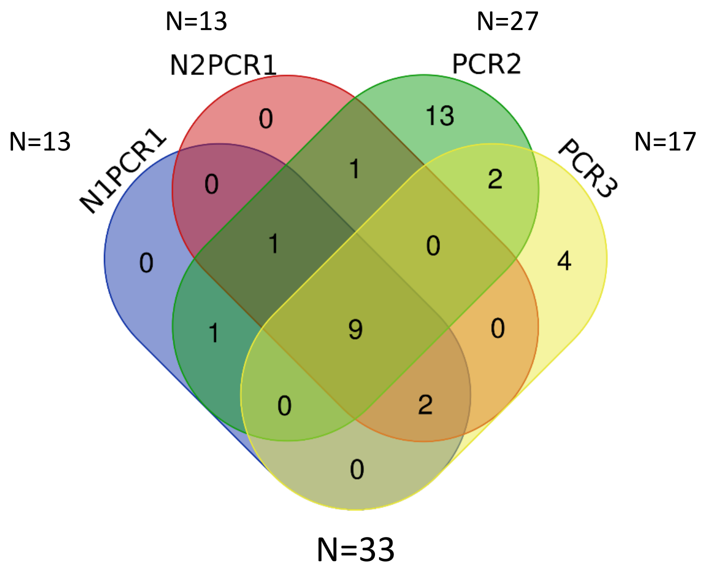


| PCRs | Total | Class |
|---|---|---|
| N1PCR1 N2PCR1 PCR2 PCR3 | 9 | Alphaproteobacteria, Fusobacteriia, Bacilli, Deinococci, Gammaproteobacteria, Actinobacteria, Acidimicrobiia, Thermoleophilia, Clostridia |
| N1PCR1 N2PCR1 PCR2 | 1 | Negativicutes |
| N1PCR1 N2PCR1 PCR3 | 2 | Longimicrobia, Bacteroidia |
| N1PCR1 PCR2 | 1 | Holophagae |
| N2PCR1 PCR2 | 1 | Coriobacteriia |
| PCR2 PCR3 | 2 | Deltaproteobacteria, Gemmatimonadetes |
| PCR2 | 13 | Gitt-GS-136, Blastocatellia (Subgroup 4), Verrucomicrobiae, Nitrospira, Oxyphotobacteria, Subgroup 6, S0134 terrestrial group, Anaerolineae, Chloroflexia, KD4-96, Babeliae, Erysipelotrichia, Planctomycetacia |
| PCR3 | 4 | Saccharimonadia, Rubrobacteria, TK10 Armatimonadia |

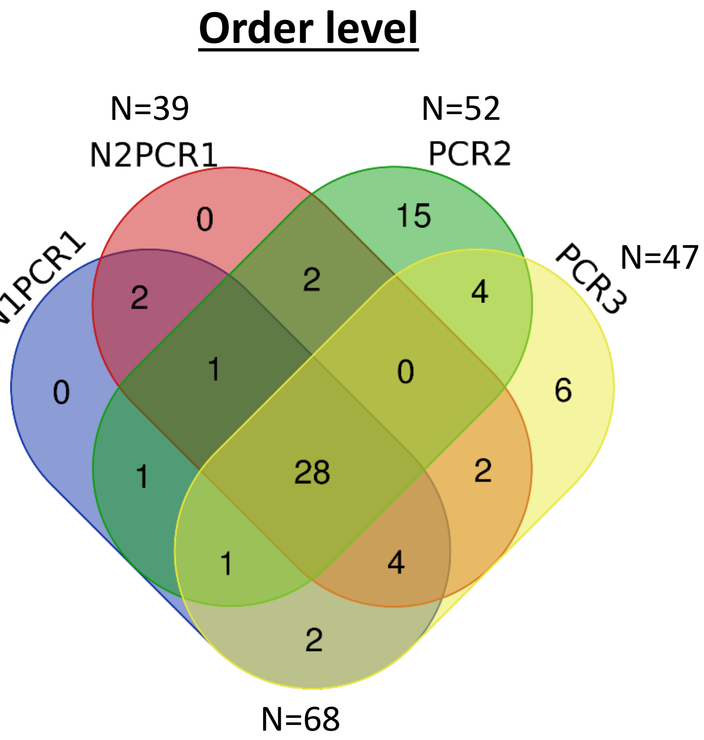


| PCRs | Total | Order |
|---|---|---|
| N1PCR1 N2PCR1 PCR2 PCR3 | 28 | Rhizobiales, Acetobacterales, Pseudonocardiales, Kineosporiales, Oceanospirillales, Xanthomonadales, Propionibacteriales, Betaproteobacteriales, Rhodobacterales, Pasteurellales, Solirubrobacterales, Caulobacterales, Frankiales, Lactobacillales, Deinococcales, Bacillales, Alteromonadales, Pseudomonadales, Fusobacteriales, Corynebacteriales, Micrococcales, Microtrichales, Actinomycetales, Vibrionales, Sphingomonadales, Enterobacteriales, Clostridiales, Thermales |
| N1PCR1 N2PCR1 PCR2 | 1 | Selenomonadales |
| N1PCR1 N2PCR1 PCR3 | 4 | Flavobacteriales, Azospirillales, Longimicrobiales, Bacteroidales |
| N1PCR1 PCR2 PCR3 | 1 | Streptomycetales |
| N1PCR1 N2PCR1 | 2 | Sneathiellales, Sphingobacteriales |
| N1PCR1 PCR2 | 1 | Subgroup 7 |
| N1PCR1 PCR3 | 2 | Cellvibrionales, Micromonosporales |
| N2PCR1 PCR2 | 2 | Aeromonadales, Coriobacteriales |
| N2PCR1 PCR3 | 2 | Cytophagales, Chitinophagales |
| PCR2 PCR3 | 4 | Gaiellales, Myxococcales, Streptosporangiales, Gemmatimonadales |
| PCR2 | 15 | Erysipelotrichales, Bdellovibrionales, Babeliales, Blastocatellales, Gemmatales, Chthoniobacterales, Salinisphaerales, SJA-15, Nitrospirales Unknown Order, Thermomicrobiales, Bifidobacteriales, Nostocales, Verrucomicrobiales, Opitutales |
| PCR3 | 6 | Rubrobacterales, Armatimonadales, Saccharimonadales, Gammaproteobacteria Incertae Sedis, Oligoflexales, Desulfovibrionales |

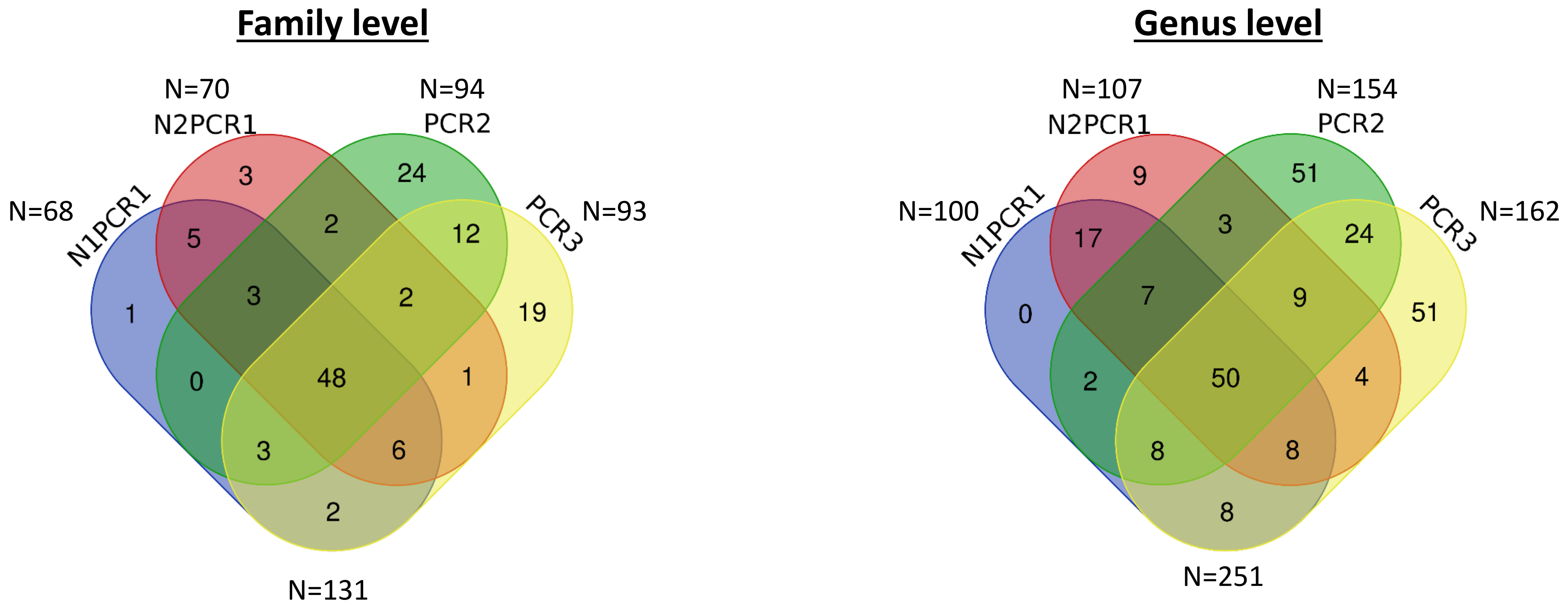
Family level
N=70
N2PCR1
N=94
PCR2
N=68
N1PCR1
PCR3
N=93
3
24
5
2
12
1
3
2
19
0
48
1
3
6
2
N=131
Genus level
N=107
N2PCR1
N=154
PCR2
N=100
N1PCR1
PCR3
N=162
9
51
17
3
24
0
7
9
51
2
50
4
8
8
8
N=251

#### Family level

| PCRs | Total | Family |
|---|---|---|
| N1PCR1 N2PCR1 PCR2 PCR3 | 48 | Intrasporangiaceae, Enterobacteriaceae, Vibrionaceae, Iamiaceae, Sanguibacteraceae, Thermaceae, Geodermatophilaceae, Neisseriaceae, Paenibacillaceae, Staphylococcaceae, Rhizobiaceae, Marinomonadaceae, Pasteurellaceae, Dermacoccaceae, Kineosporiaceae, Planococcaceae, Microbacteriaceae, Micrococcaceae, Rhodobacteraceae, Bacillaceae, Corynebacteriaceae, Streptococcaceae, Nocardioidaceae, Pseudomonadaceae, Fusobacteriaceae, Carnobacteriaceae, Brevibacteriaceae, Family, XI, Solirubrobacteraceae, Leptotrichiaceae, Pseudonocardiaceae, Beijerinckiaceae, Xanthobacteraceae, Enterococcaceae, Listeriaceae, Deinococcaceae, Actinomycetaceae, Moraxellaceae, Lachnospiraceae, Cellulomonadaceae, Acetobacteraceae, Ilumatobacteraceae, Lactobacillaceae, Xanthomonadaceae, Burkholderiaceae, Caulobacteraceae, Sphingomonadaceae, Propionibacteriaceae |
| N1PCR1 N2PCR1 PCR2 | 3 | Promicromonosporaceae, Nocardiaceae, Veillonellaceae |
| N1PCR1 N2PCR1 PCR3 | 6 | Prevotellaceae, Dermabacteraceae, Weeksellaceae, Longimicrobiaceae, Azospirillaceae, Flavobacteriaceae |
| N1PCR1 PCR2 PCR3 | 3 | Rhodanobacteraceae, Streptomycetaceae, Dermatophilaceae |
| N2PCR1 PCR2 PCR3 | 2 | Sporolactobacillaceae, Ruminococcaceae |
| N1PCR1 N2PCR1 | 5 | Sporichthyaceae, Shewanellaceae, Sneathiellaceae, Porphyromonadaceae, Methylophilaceae |
| N1PCR1 PCR3 | 2 | Cellvibrionaceae, Micromonosporaceae |
| N2PCR1 PCR2 | 2 | Aeromonadaceae, Leuconostocaceae |
| N2PCR1 PCR3 | 1 | Chitinophagaceae |
| PCR2 PCR3 | 12 | Mycobacteriaceae, 67-14, Clostridiaceae, 1, Aerococcaceae, Unknown, Family, Nocardiopsaceae, Gaiellaceae, Streptosporangiaceae, Alteromonadaceae, Idiomarinaceae, Gemmatimonadaceae, Hyphomicrobiaceae |
| N1PCR1 | 1 | NS11-12 marine group |
| N2PCR1 | 3 | Spirosomaceae Sphingobacteriaceae Coriobacteriaceae |
| PCR2 | 24 | Solimonadaceae, Rhodocyclaceae, Nostocaceae, Bifidobacteriaceae, Blastocatellaceae, Chroococcidiopsaceae, Chthoniobacteraceae, Psychromonadaceae, Thermoactinomycetaceae, Rubritaleaceae, Peptostreptococcaceae, Frankiaceae, Nitrospiraceae, Family, XIII, Opitutaceae, Bdellovibrionaceae, Atopobiaceae, Family, XVII, Myxococcaceae, Family, X, Erysipelotrichaceae, Vermiphilaceae, Gemmataceae, JG30-KF-CM45 |
| PCR3 | 19 | Bacteroidaceae, 0319-6G20, BIrii41, Devosiaceae, Trueperaceae, Dysgonomonadaceae, Hymenobacteraceae, Desulfovibrionaceae, Nitrosomonadaceae, Rubrobacteriaceae, Halomonadaceae, Jonesiaceae, Demequinaceae, Phaselicystidaceae, Microtrichaceae, Saccharimonadaceae, Haliangiaceae, Prolixibacteraceae, Nakamurellaceae |

### Genus level

| PCRs | Total | Genus |
|---|---|---|
| N1PCR1 N2PCR1 PCR2 PCR3 | 50 | Schlegelella, Turicella, Listeria, Actinomyces, Psychrobacter, Brevibacillus, Arthrobacter, Brochothrix, Lawsonella, Peptoniphilus, Brevibacterium, Vibrio, Massilia, Enterococcus, Anaerococcus, Ralstonia, Leptotrichia, Fusobacterium, Dolosigranulum, Gemella, Meiothermus, Rothia, Haemophilus, Novosphingobium, Pectobacterium, Marinomonas, Paracoccus, Sphingomonas, Dermacoccus, Cutibacterium, Bradyrhizobium, Cellulomonas, Iamia, Staphylococcus, Corynebacterium, Deinococcus, Lactobacillus, Microvirga, Roseomonas, Nocardioides, Streptococcus, Finegoldia, Blastococcus, Pseudomonas, Bacillus, Micrococcus, Pseudonocardia, Stenotrophomonas, Sanguibacter, Kocuria |
| N1PCR1 N2PCR1 PCR2 | 7 | Brevundimonas, Variovorax, Rubellimicrobium, Jeotgalicoccus, Alloiococcus, Acidovorax, Carnobacterium |
| N1PCR1 N2PCR1 PCR3 | 8 | Brachybacterium, Solirubrobacter, Skermanella, Alloprevotella, Frigoribacterium, Prevotella, Ornithinimicrobium, Chryseobacterium |
| N1PCR1 PCR2 PCR3 | 8 | Streptomyces, Microbacterium, Afipia, Jiangella, Modestobacter, Diaphorobacter, Neisseria, Methylobacterium |
| N2PCR1 PCR2 PCR3 | 9 | Allorhizobium-Neorhizobium-Pararhizobium-Rhizobium, Bosea, Paenibacillus, Salmonella, Phenylobacterium, Escherichia-Shigella, Pantoea, Acinetobacter, Granulicatella |
| N1PCR1 N2PCR1 | 17 | Capnocytophaga, hgcI clade, Burkholderia-Caballeronia-Paraburkholderia, Photobacterium, Porphyromonas, Ensifer, Citricoccus, Pseudokineococcus, Lautropia, Oligella, Macrococcus, Pseudoxanthomonas, Gordonia, Ferrovibrio, Shewanella, Dialister, Promicromonospora |
| N1PCR1 PCR2 | 2 | Mesorhizobium, Aquipuribacter |
| N1PCR1 PCR3 | 8 | Geodermatophilus, Empedobacter, Rudaea, Cellvibrio, Lysinibacillus, Fusicatenibacter, Cloacibacterium, Flavobacterium |
| N2PCR1 PCR2 | 3 | Leuconostoc, Aeromonas, Serratia |
| N2PCR1 PCR3 | 4 | Domibacillus, Ammoniphilus, Quadrisphaera, Microlunatus |
| PCR2 PCR3 | 24 | Tepidimonas, Anoxybacillus, Propioniciclava, Aerococcus, Pseudarthrobacter, Agrococcus, Varibaculum, Ramlibacter, Clostridium, sensu, stricto, Rheinheimera, Nocardiopsis, Enterobacter, Nonomuraea, Actinotalea, Parvimonas, Gaiella, Saccharopolyspora, Lysobacter, Hyphomicrobium, Mycobacterium, Idiomarina, Curtobacterium, Aquabacterium, Naasia |
| N2PCR1 | 9 | Pedobacter, Collinsella, Aggregatibacter, Knoellia, Craurococcus, Leifsonia, Dyadobacter, Enhydrobacter, Flavisolibacter |
| PCR2 | 51 | Altererythrobacter, Undibacterium, Stenotrophobacter, Bifidobacterium, Jatrophihabitans, Candidatus, Udaeobacter, Actinobacillus, Psychromonas, Niveibacterium, Gemmatimonas, Rhodoferax, Nitrospira, Luteolibacter, Bdellovibrio, Limnohabitans, Rhodococcus, Noviherbaspirillum, Turicibacter, Nevskia, [Eubacterium], brachy, group, Lechevalieria, Morganella, Luteitalea, Oxalobacter, [Eubacterium], yurii, group, Ethanoligenens, Amaricoccus, Pseudoclavibacter, Thermaerobacter, Aliterella, CENA595, Marmoricola, Kingella, Thermicanus, Sphingopyxis, S31, Calothrix, PCC-6303, Blautia, Kluyvera, Novibacillus, Opitutus, Veillonella, Weissella, Fimbriiglobus, Rhizobacter, Kurthia, Kineococcus, Tuberibacillus, Prauserella, Tetrasphaera, Roseococcus, Atopobium |
| PCR3 | 51 | Glutamicibacter, Gemmobacter, Vulcaniibacterium, Bacteroides, Halomonas, Ezakiella, Caulobacter, Adhaeribacter, Ignavigranum, MND1, Faecalibacterium, Alishewanella, Phaselicystis, Aneurinibacillus, Oceanobacillus, Dermabacter, Bergeyella, Porphyrobacter, Luteimonas, Erythrobacter, Actinoplanes, Hymenobacter, Flavihumibacter, Abiotrophia, Sphingoaurantiacus, Pseudorhodoferax, Tatumella, Desulfovibrio, Nakamurella, Rubrobacter, Chryseomicrobium, Microcella, Devosia, Proteiniclasticum, Belnapia, Thermomonas, Leptothrix, Jonesia, Proteiniphilum, Roseburia, Methylocella, Qipengyuania, Conyzicola, metagenome, WCHB1-32, Haliangium, Friedmanniella, Truepera, Lysinimicrobium, Acidibacter, Lachnoanaerobaculum |
