## Supplementary figures and images for "Evaluation of established methods for DNA extraction and primer pairs targeting 16S rRNA gene for bacterial microbiome profiling of olive xylem sap"

### Suppl Figure S1

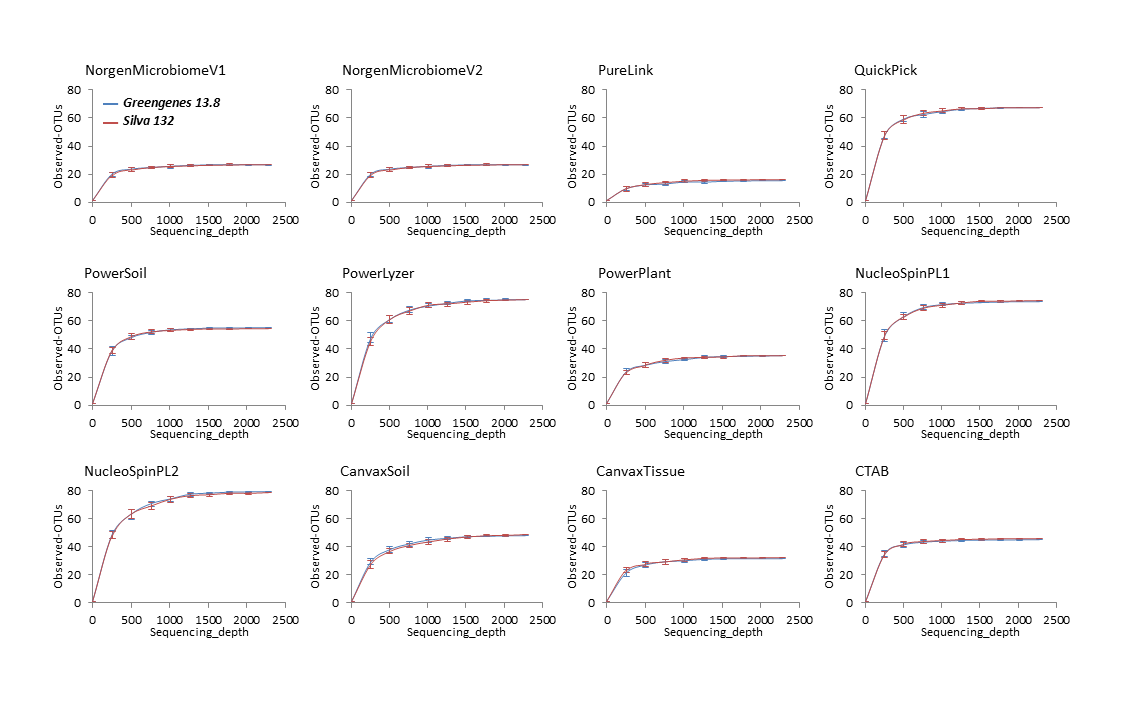
