## Supplementary material for "Evaluation of established methods for DNA extraction and primer pairs targeting 16S rRNA gene for bacterial microbiome profiling of olive xylem sap": Suppl Figure S2

### Phylum level

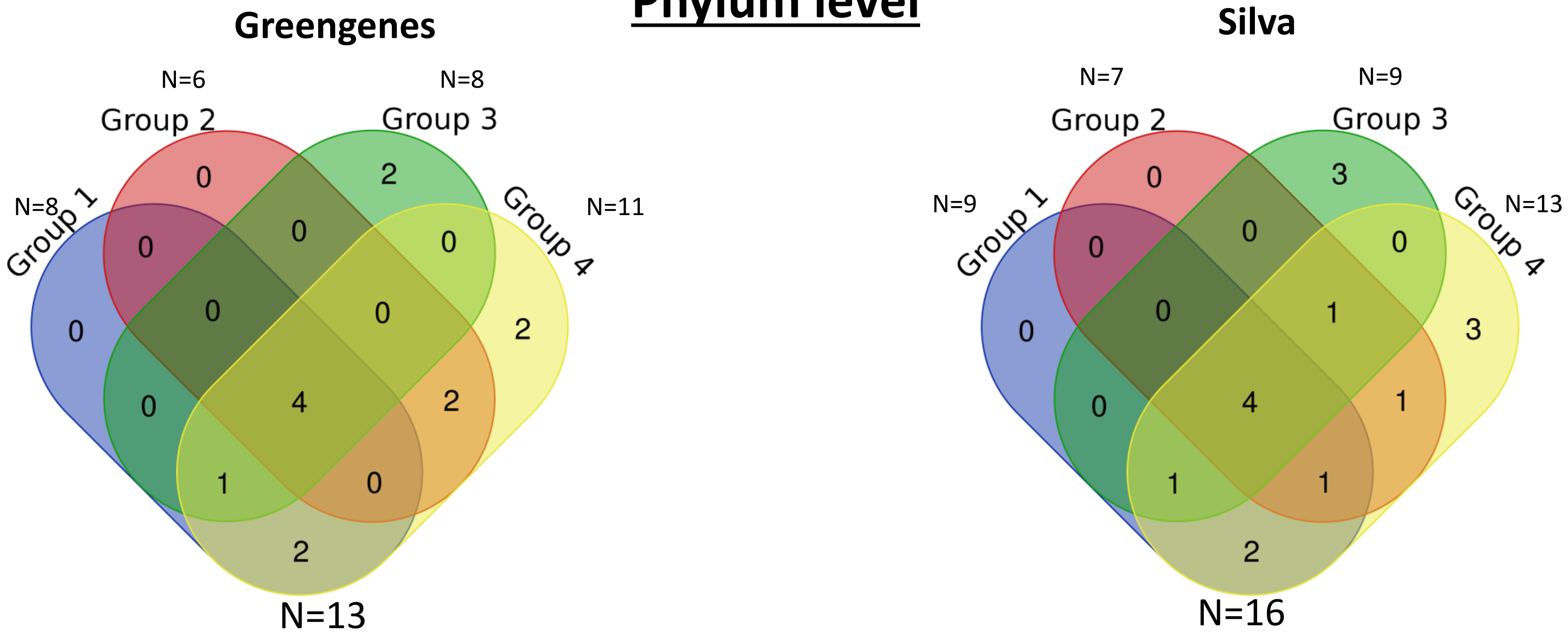


| DNA extraction kits | Total | Phylum (Greengenes) | Phylum (Silva) |
|---|---|---|---|
| Group 1 Group 2 Group 3 Group 4 | 5 | Actinobacteria, Proteobacteria, Bacteroidetes, Firmicutes | Bacteroidetes, Actinobacteria, Proteobacteria, Firmicutes |
| Group 1 Group 2 Group 4 | 0/1 | | D_0_Bacteria |
| Group 1 Group 3 Group 4 | 1 | Acidobacteria | Acidobacteria |
| Group 2 Group 3 Group 4 | 0/1 | | Cyanobacteria |
| Group 1 Group 4 | 2 | [Thermi], Planctomycetes | Planctomycetes, Deinococcus-Thermus |
| Group 2 Group 4 | 1 | Fusobacteria | Fusobacteria |
| Group 3 | 2/3 | Chloroflexi, Nitrospirae | Nitrospirae, Fibrobacteres, Chloroflexi |
| Group 4 | 2/3 | Verrucomicrobia, Gemmatimonadetes | Saccharibacteria, Gemmatimonadetes, Verrucomicrobia |

### Class level

#### Greengenes

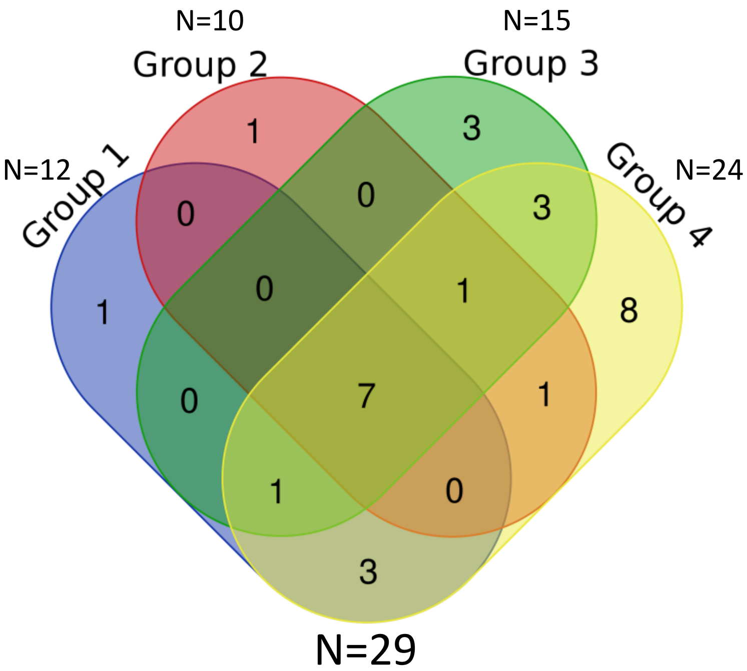


#### Silva

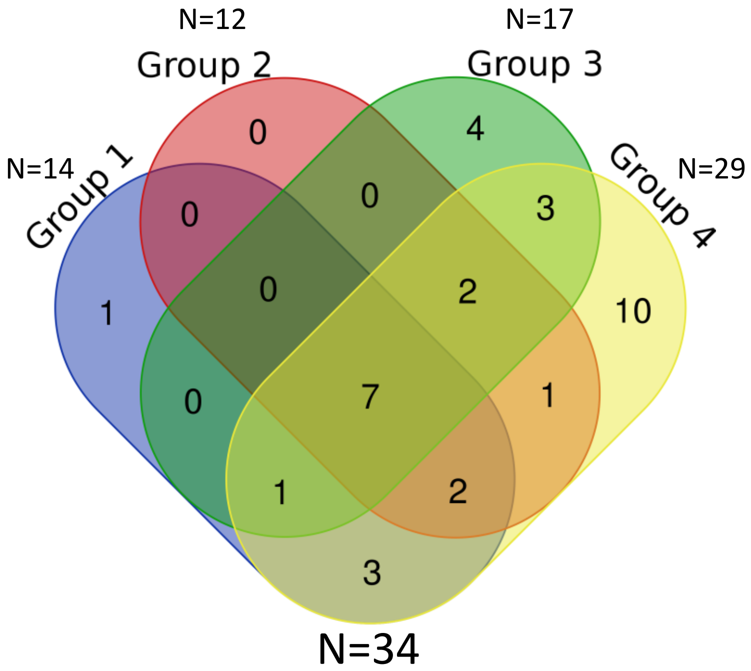


| DNA extraction kits | Total | Class (Greengenes) | Class (Silva) |
|---|---|---|---|
| Group 1 Group 2 Group 3 Group 4 | 7 | Alphaproteobacteria, [Saprospirae], Betaproteobacteria, Actinobacteria, Clostridia, Bacilli, Gammaproteobacteria | Alphaproteobacteria, Sphingobacteriia, Bacilli, Gammaproteobacteria, Actinobacteria, Betaproteobacteria, Clostridia |
| Group 1 Group 2 Group 4 | 0/2 | | Negativicutes, D_0_Bacteria |
| Group 1 Group 3 Group 4 | 1 | Deltaproteobacteria | Deltaproteobacteria |
| Group 2 Group 3 Group 4 | 1/2 | Bacteroidia | Cyanobacteria, Bacteroidia |
| Group 1 Group 4 | 3 | Deinococci, Solibacteres, Planctomycetia | Deinococci, Solibacteres, Planctomycetacia |
| Group 2 Group 4 | 1 | Fusobacteriia | Fusobacteriia |
| Group 3 Group 4 | 3 | Acidimicrobiia, Cytophagia, Flavobacteriia | Flavobacteriia, Cytophagia, Acidimicrobiia |
| Group 1 | 1 | DA052 | Subgroup 2 |
| Group 2 | 1/0 | Synechococcophycideae | |
| Group 3 | 3/4 | Nitrospira, Anaerolineae, Holophagae | Holophagae, Nitrospira, Fibrobacteria, Anaerolineae |
| Group 4 | 8/10 | Gemm-3, Verrucomicrobiae, Erysipelotrichi, Thermoleophilia, [Spartobacteria], OM190, Oscillatoriophycideae, [Rhodothermi] | Bacteroidetes Incertae Sedis, OM190, Verrucomicrobiae D_0_Bacteria;Saccharibacteria, Longimicrobia; Spartobacteria; Gemmatimonadetes; Thermoleophilia, Erysipelotrichia, D_0_Bacteria;Ambiguous_taxa |

### Order level

#### Greengenes

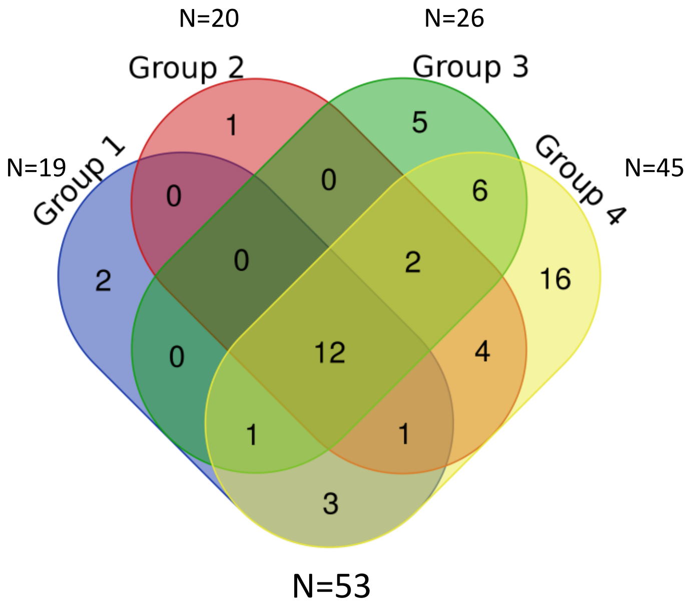


#### Silva

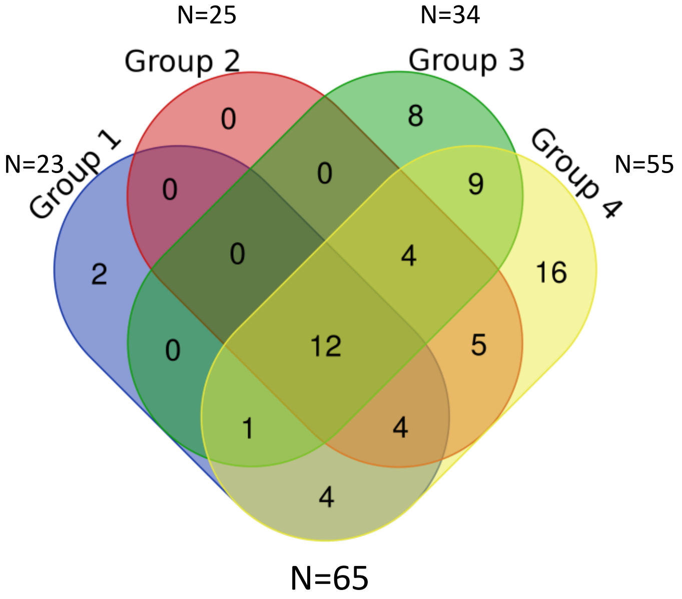


| DNA extraction kits | Total | Order (Greengenes) | Order (Silva) |
|---|---|---|---|
| Group 1 Group 2 Group 3 Group 4 | 12 | Bacillales, Pseudomonadales, Sphingomonadales, Xanthomonadales, [Saprospirales], Alteromonadales, Rhizobiales, Clostridiales, Actinomycetales, Gemellales, Burkholderiales, Lactobacillales | Rhizobiales, Xanthomonadales, Propionibacteriales, Lactobacillales, Burkholderiales, Bacillales, Pseudomonadales, Corynebacteriales, Micrococcales, Sphingobacteriales, Sphingomonadales, Clostridiales |
| Group 1 Group 2 Group 4 | 1/4 | Enterobacteriales | Selenomonadales, Chromatiales, D_0_Bacteria, Enterobacteriales |
| Group 1 Group 3 Group 4 | 1 | Rhodospirillales | Rhodospirillales |
| Group 2 Group 3 Group 4 | 2/4 | Bacteroidales, Caulobacterales | Rhodobacterales, Frankiales, Caulobacterales, Bacteroidales |
| Group 1 Group 4 | 3/4 | Deinococcales, Myxococcales, Solibacterales | Pseudonocardiales, Planctomycetales, Deinococcales, Solibacterales |
| Group 2 Group 4 | 4/5 | Vibrionales, Neisseriales, Pasteurellales, Fusobacteriales | Neisseriales, SubsectionIII, Pasteurellales, Fusobacteriales, Vibrionales |
| Group 3 Group 4 | 6/9 | Methylophilales, Rhodocyclales, Rhodobacterales, Flavobacteriales, Cytophagales, Acidimicrobiales | Cytophagales, Kineosporiales, Rickettsiales, SubsectionII, Methylophilales, Actinomycetales, Rhodocyclales, Acidimicrobiales, Flavobacteriales |
| Group 1 | 2 | Ellin6513, Gemmatales | D_2_Subgroup 2, Oligoflexales |
| Group 2 | 1/0 | Pseudanabaenales | |
| Group 3 | 5/8 | Nitrospirales, Holophagales, Bdellovibrionales, c_Betaproteobacteria, SBR1031 | Cellvibrionales, Holophagales, Bdellovibrionales, PeM15, Fibrobacterales, Nitrospirales, Nitrosomonadales, Anaerolineales |
| Group 4 | 16/16 | Ellin329, Chroococcales, Solirubrobacterales, Aeromonadales, Procabacteriales, Oscillatoriales, agg27, Thermales, [Chthoniobacterales], Planctomycetales, Rickettsiales, [Rhodothermales], Verrucomicrobiales, Gemm-3, Erysipelotrichales, Oceanospirillales | Erysipelotrichales, Aeromonadales, Oceanospirillales, Chthoniobacterales, D_2_Betaproteobacteria, Solirubrobacterales, D_0_Bacteria; Ambiguous_taxa, D_2_OM190, Myxococcales, Verrucomicrobiales, Streptosporangiales, Gemmatimonadales, D_1_Saccharibacteria, Longimicrobiales, Order II, Thermales |

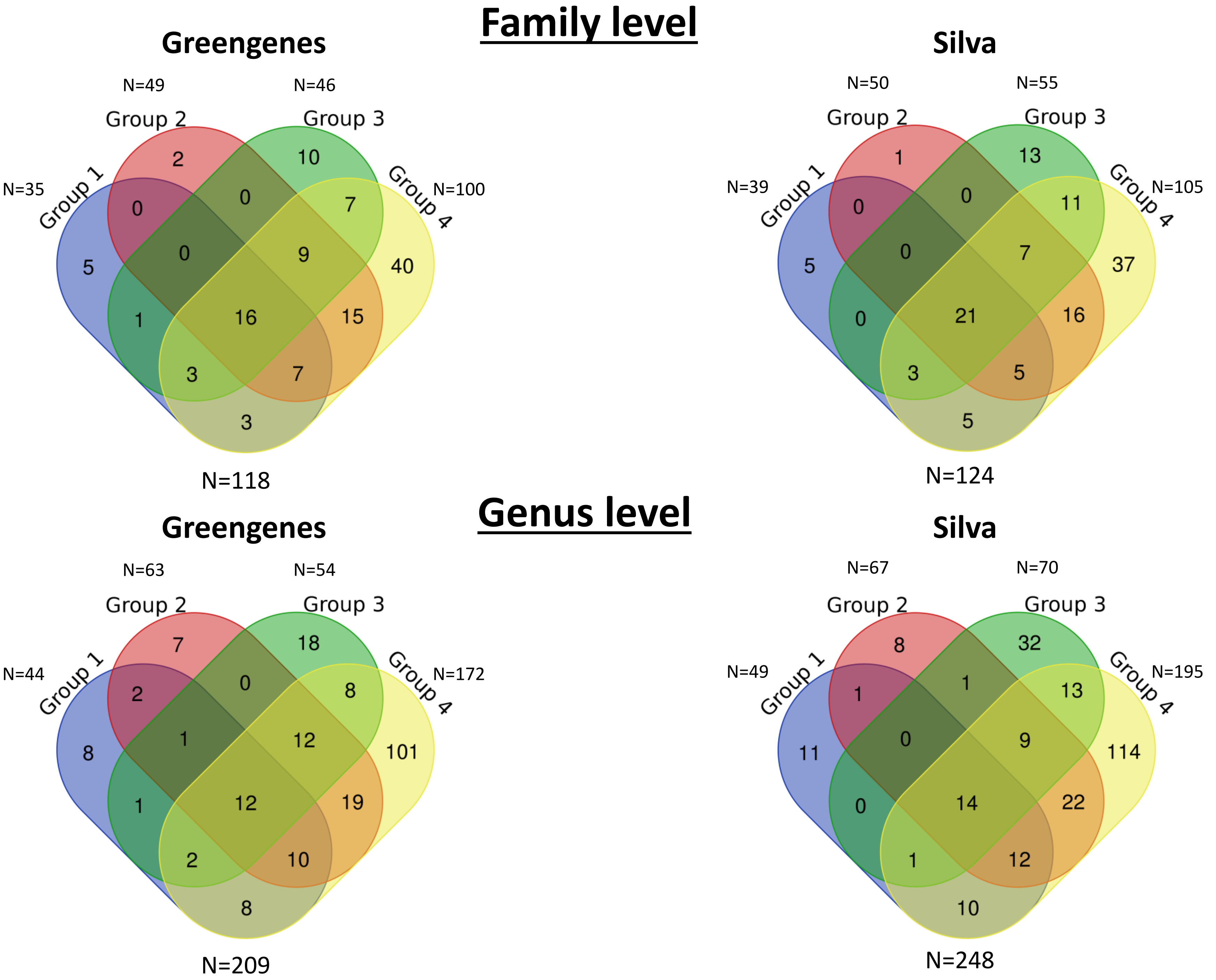

Family level
Greengenes
N=49
N=46
Group 2
Group 3
N=35
Group 1
Group 4
N=100
2
10
0
0
7
5
0
9
40
1
16
15
3
7
3
N=118
Silva
N=50
N=55
Group 2
Group 3
N=39
Group 1
Group 4
N=105
1
13
0
0
11
5
0
7
37
0
21
16
3
5
5
N=124
Genus level
Greengenes
N=63
N=54
Group 2
Group 3
N=44
Group 1
Group 4
N=172
7
18
2
0
8
8
1
12
101
1
12
19
2
10
8
N=209
Silva
N=67
N=70
Group 2
Group 3
N=49
Group 1
Group 4
N=195
8
32
1
1
13
11
0
9
114
0
14
22
1
12
10
N=248

### Family level

| DNA extraction kits | Total | Family (Greengenes) | Family (Silva) |
|---|---|---|---|
| Group 1 Group 2 Group 3 Group 4 | 16/21 | Corynebacteriaceae, Comamonadaceae, Staphylococcaceae, Chitinophagaceae, Pseudomonadaceae, Lactobacillaceae, Methylobacteriaceae, Streptococcaceae, Rhizobiaceae, Xanthomonadaceae, Gemellaceae, Propionibacteriaceae, [Tissierellaceae], Oxalobacteraceae, Microbacteriaceae, Micrococcaceae | Xanthobacteraceae, Bradyrhizobiaceae, Oxalobacteraceae, Staphylococcaceae, Rhizobiaceae, Comamonadaceae, Family XI (Bacillales), Microbacteriaceae, Micrococcaceae, Corynebacteriaceae, Methylobacteriaceae, Lactobacillaceae, Streptococcaceae, Chitinophagaceae, Family XI (Clostridiales), Pseudomonadaceae, Xanthomonadaceae, Burkholderiaceae, Carnobacteriaceae, Sphingomonadaceae, Propionibacteriaceae |
| Group 1 Group 2 Group 4 | 7/5 | Burkholderiaceae, Veillonellaceae, [Chromatiaceae], Bradyrhizobiaceae, Enterobacteriaceae, Sphingomonadaceae, o_Lactobacillales | Enterobacteriaceae, Chromatiaceae, Veillonellaceae, D_0_Bacteria, , D_3_Sphingomonadales |
| Group 1 Group 3 Group 4 | 3/3 | Lachnospiraceae, Alcaligenaceae, Rhodospirillaceae | Lachnospiraceae, Alcaligenaceae, Rhodospirillales Incertae Sedis |
| Group 2 Group 3 Group 4 | 9/7 | o_Actinomycetales, Nocardioidaceae, ACK-M1, Caulobacteraceae, Hyphomicrobiaceae, [Paraprevotellaceae], o_Rhizobiales, Listeriaceae, Phyllobacteriaceae | Sporichthyaceae, Phyllobacteriaceae, Listeriaceae, Rhodobacteraceae, Prevotellaceae, Nocardioidaceae, Caulobacteraceae |
| Group 1 Group 3 | 1/0 | Sinobacteraceae | |
| Group 1 Group 4 | 3/5 | Pseudonocardiaceae, o_Solibacterales, Deinococcaceae | Pseudonocardiaceae, Planctomycetaceae, Deinococcaceae, Solibacteraceae (Subgroup 3), I-10 |
| Group 2 Group 4 | 15/16 | Neisseriaceae, Mycobacteriaceae, Vibrionaceae, Fusobacteriaceae, Paenibacillaceae, Pasteurellaceae, Erythrobacteraceae, Bacillaceae, Moraxellaceae, Ruminococcaceae, Intrasporangiaceae, Porphyromonadaceae, Aerococcaceae, Prevotellaceae, Dermabacteraceae | Intrasporangiaceae, Mycobacteriaceae, Vibrionaceae, FamilyI, Neisseriaceae, Paenibacillaceae, Pasteurellaceae, Dermacoccaceae, Moraxellaceae, Bacillaceae, Porphyromonadaceae, Ruminococcaceae, Aerococcaceae, Erythrobacteraceae, Fusobacteriaceae, Dermabacteraceae |
| Group 3 Group 4 | 7/11 | Rhodocyclaceae, Cytophagaceae, Methylophilaceae, Nocardiaceae, Actinomycetaceae, Rhodobacteraceae, Flavobacteriaceae | Rhodocyclaceae, Nocardiaceae, Actinomycetaceae, D_3_Frankiales, Kineosporiaceae, Methylophilaceae, Rhizobiales Incertae Sedis, FamilyII, Cytophagaceae, Hyphomicrobiaceae, Flavobacteriaceae |
| Group 1 | 5/5 | 0319-6G20, Methylocystaceae, o_Ellin6513, Xanthobacteraceae, Gemmataceae | 0319-6G20, D_2_Subgroup 2, Methylocystaceae, Xanthomonadales Incertae Sedis, Nevskiaceae |
| Group 2 | 2/1 | Brevibacteriaceae, Pseudanabaenaceae | Brevibacteriaceae |
| Group 3 | 10/13 | Carnobacteriaceae, Nitrospiraceae, c_Betaproteobacteria, o_Sphingomonadales, A4b, Microthrixaceae, Alteromonadaceae, Holophagaceae, Bdellovibrionaceae, Hyphomonadaceae | Rickettsiales Incertae Sedis, Cryptosporangiaceae, Bdellovibrionaceae, D_3_Xanthomonadales, D_3_PeM15, Anaerolineaceae, Fibrobacteraceae, Hyphomonadaceae, Cellvibrionaceae, Nitrospiraceae, Acidimicrobiales Incertae Sedis, Nitrosomonadaceae, Holophagaceae |
| Group 4 | 40/37 | Thermoactinomycetaceae, Verrucomicrobiaceae, Sporichthyaceae, Procabacteriaceae, Peptostreptococcaceae, Erysipelotrichaceae, Acetobacteraceae, o_Solirubrobacterales, Haliangiaceae, Gordoniaceae, Rhodothermaceae, Planococcaceae, Xenococcaceae, Leuconostocaceae, o_Rickettsiales, Nocardiopsaceae, o_Myxococcales, Aeromonadaceae, Geodermatophilaceae, Phormidiaceae, Oceanospirillaceae, o_Ellin329, [Exiguobacteraceae], Bacillales, Clostridiaceae, C111, [Chthoniobacteraceae], o_agg27, [Weeksellaceae], Rickettsiaceae, Trueperaceae, [Mogibacteriaceae], Planctomycetaceae, o_Clostridiales, Cellulomonadaceae, c_Gemm-3, Peptococcaceae, Conexibacteraceae, Thermaceae, Williamsiaceae | BIrii41, Clostridiales vadinBB60 group, Trueperaceae, Thermaceae, Rhodothermaceae, Acidimicrobiaceae, Rickettsiaceae, Nocardiopsaceae, Geodermatophilaceae, Sandaracinaceae, Myxococcaceae, D_2_Betaproteobacteria, Chthoniobacteraceae, Thermoactinomycetaceae, Longimicrobiaceae, Family XII, Erysipelotrichaceae, Elev-16S-1332, D_2_OM190, Peptostreptococcaceae, Planococcaceae, D_0_Bacteria; Ambiguous_taxa, Cellulomonadaceae, Acetobacteraceae, Christensenellaceae, Clostridiaceae 1, Peptococcaceae, Haliangiaceae, Gemmatimonadaceae, D_1_Saccharibacteria, Aeromonadaceae, Leuconostocaceae, Verrucomicrobiaceae, Family XIII, Rhodospirillaceae, YNPFFP1, Oceanospirillaceae |

### Genus level

| DNA extraction kits | Total | Genus (Greengenes) | Genus (Silva) |
|---|---|---|---|
| Group 1 Group 2 Group 3 Group 4 | 12/14 | f_Comamonadaceae, Pseudomonas, Sediminibacterium f_Oxalobacteraceae, Streptococcus, Propionibacterium, Microbacterium, Agrobacterium, Lactobacillus, Corynebacterium, Gemellaceae, Staphylococcus | Corynebacterium 1, Microbacterium, Lawsonella, D_4_Comamonadaceae, Rhizobium, Gemella, Propionibacterium, Staphylococcus, Lactobacillus, Streptococcus, Aquabacterium, Granulicatella, Methylobacterium, Pseudomonas |
| Group 1 Group 2 Group 3 | 1/0 | f_Methylobacteriaceae | |
| Group 1 Group 2 Group 4 | 10/12 | Stenotrophomonas, Veillonella, f_Enterobacteriaceae, f_Sphingomonadaceae, Rheinheimera, Anaerococcus, Ralstonia, Bradyrhizobiaceae, Afipia, o_Lactobacillales | Massilia, Anaerococcus, Ralstonia, Rheinheimera, D_4_Chitinophagaceae, D_0_Bacteria, Pantoea, Sphingomonas, Bradyrhizobium, Veillonella, D_3_Sphingomonadales, Stenotrophomonas |
| Group 1 Group 3 Group 4 | 2/1 | f_Xanthomonadaceae, f_Rhodospirillaceae | Massilia |
| Group 2 Group 3 Group 4 | 12/9 | o_Actinomycetales, f_Nocardioidaceae, f_ACK-M1, Finegoldia, Rhodoplanes, Brochothrix, o_Rhizobiales, f_Phyllobacteriaceae, Mycoplana, [Prevotella], f_Microbacteriaceae, Kocuria | Bosea, Brochothrix, Brevundimonas, hgcI clade, Alloprevotella, Paracoccus, Mesorhizobium, Finegoldia, Kocuria |
| Group 1 Group 2 | 2/1 | Burkholderia, Bradyrhizobium | Burkholderia-Paraburkholderia |
| Group 1 Group 3 | 1/0 | f_Sinobacteraceae | |
| Group 1 Group 4 | 8/10 | Achromobacter, Pseudonocardia, Micrococcus, Delftia,o_Solibacterales, Deinococcus, Novosphingobium, Janthinobacterium | Delftia, Achromobacter, Escherichia-Shigella, Enterobacter, Novosphingobium, Deinococcus, Bryobacter, D_4_I-10, Micrococcus, Pseudonocardia |
| Group 2 Group 3 | 0/1 | | Variibacter |
| Group 2 Group 4 | 19/22 | Bacillus, Mycobacterium, Neisseria, f_Erythrobacteraceae, Diaphorobacter, Peptoniphilus, f_Aerococcaceae, Paenibacillus, Fusobacterium, Haemophilus, Porphyromonas, Methylobacterium, Prevotella, f_Intrasporangiaceae, Acinetobacter, Aerococcus, Anoxybacillus, Rothia, Photobacterium | D_4_Intrasporangiaceae, D_4_Enterobacteriaceae, D_4_Vibrionaceae, Anoxybacillus, Paenibacillus, Aerococcus, Peptoniphilus, Porphyromonas, Fusobacterium, Porphyrobacter, Rothia, Haemophilus, Abiotrophia, Dermacoccus, Prevotella, Pseudoclavibacter, Diaphorobacter, Acinetobacter, Neisseria, Nocardioides, Mycobacterium, Bacillus |
| Group 3 Group 4 | 8/13 | Hyphomicrobium, f_Lachnospiraceae, Paracoccus, f_Methylophilaceae, Flavobacterium, Actinomyces, Limnohabitans, Rhodococcus | Actinomyces, D_3_Frankiales, Methylotenera, Rubellimicrobium, D_4_Microbacteriaceae, Limnohabitans, Rhodococcus, Rhodobacter, Marmoricola, Hyphomicrobium, D_4_Xanthomonadaceae, Flavobacterium, D_4_Sphingomonadaceae |
| Group 1 | 8/11 | f_0319-6G20, Roseburia, o_Ellin6513, Labrys, Gemmata, Actinomycetospora, Nevskia, Methylosinus | D_4_0319-6G20, Labrys, Duganella, Methylosinus, Luteibacter, Gemmata, D_2_Subgroup 2, Nevskia, Actinomycetospora, Roseburia, Acidibacter |
| Group 2 | 7/8 | Dermabacter, Moraxella, Klebsiella, f_Ruminococcaceae, Brevibacterium, Sphingomonas, Leptolyngbya | D_4_Bradyrhizobiaceae, Leptolyngbya, Brevibacterium, Moraxella, Knoellia, D_4_Dermabacteraceae, Klebsiella, [Eubacterium] coprostanoligenes group |
| Group 3 | 18/32 | Cellvibrio, Candidatus Rhodoluna, Dechloromonas, Bdellovibrio Methylibium, c_Betaproteobacteria, f_Chitinophagaceae, Candidatus Aquiluna, o_Sphingomonadales, Granulicatella, f_Alcaligenaceae, f_Cytophagaceae, f_A4b, f_Microthrixaceae, Nitrospira, Methylotenera, f_Holophagaceae, f_Hyphomonadaceae | PRD01a011B, D_4_Rickettsiales Incertae Sedis, Undibacterium, Polynucleobacter, Sediminibacterium, Candidatus Aquiluna, Pleurocapsa, Woodsholea, Phreatobacter, Nitrospira, Dechloromonas, D_4_Fibrobacteraceae, Fodinicola, Bdellovibrio, Pseudokineococcus, D_4_Nitrosomonadaceae, Cellvibrio, GKS98 freshwater group, Niastella, D_3_Xanthomonadales, D_3_PeM15, Lachnoclostridium 5, Pseudarcicella, D_4_Anaerolineaceae, Methylophilus, Holophaga, Candidatus Microthrix, D_4_Methylophilaceae, Rhizobacter, Candidatus Planktoluna, OM43 clade, Candidatus Rhodoluna |
| Group 4 | 101/114 | Rubricoccus, Williamsia, f_Peptostreptococcaceae, f_Neisseriaceae, Faecalibacterium, f_Haliangiaceae, Arthrobacter, Nocardioides f_Pseudomonadaceae, f_Planococcaceae, Flavisolibacter, Exiguobacterium, f_Aeromonadaceae, f_Geodermatophilaceae, Clostridium, f_[Chromatiaceae], Hymenobacter, Parasegitibacter, Saccharopolyspora, o_Ellin329, Ruminococcus, o_Bacillales, Mogibacterium, Virgibacillus, Curtobacterium, f_C111, Selenomonas, Coprococcus, Tepidimonas, f_Propionibacteriaceae, Planomicrobium, Adhaeribacter, Erwinia, Aggregatibacter, o_Clostridiales, Chryseobacterium, Skermanella, c_Gemm-3, Marinobacterium, Blautia, Kaistobacter, Alloiococcus, f_Thermoactinomycetaceae, f_Sporichthyaceae, Chthoniobacter, f_Procabacteriaceae, Vibrio, f_Acetobacteraceae, o_Solirubrobacterales, f_Rhodocyclaceae, Janibacter, Gordonia, Prauseria, Brachybacterium, f_Xenococcaceae, Brevundimonas, Brevibacillus, Capnocytophaga, f_Leuconostocaceae, o_Rickettsiales, Planctomyces, f_Nocardiopsaceae, Marinomonas, Planktothrix, o_Myxococcales, Luteolibacter, Rickettsia, Balneimonas, Microbispora, Truepera, f_Moraxellaceae, Parvimonas, Lautropia, Actinotalea, f_Caulobacteraceae, Enhydrobacter, Agrococcus, f_Clostridiaceae, Megamonas, Geobacillus, Kingella, Peptostreptococcus, Catonella, o_agg27, f_[Weeksellaceae], Pseudoclavibacter, Bulleidia, Meiothermus, Comamonas, Asticcacaulis, Leucobacter, Peptococcus, f_Rhodobacteraceae, Aeromicrobium, f_Micrococcaceae, Deefgea, f_Conexibacteraceae, Limnobacter, Gemella, Rhodobacter, Propionicimonas | Capnocytophaga, Arenimonas, D_4_Rhodocyclaceae, Glutamicibacter, Brachybacterium, D_4_BIrii41, Aestuariimicrobium, Atopostipes, Parasegetibacter, Virgibacillus, Johnsonella, Geobacillus, Prevotella 2, Tepidimonas, Clostridium sensu stricto 1, Brevibacillus, Selenomonas 3, Rhizomicrobium, Comamonas, Catonella, Arthrobacter, Frondihabitans, D_4_Neisseriaceae, Propioniciclava, Prevotella 7, Adhaeribacter, Pseudarthrobacter, CL500-29 marine group, Peptococcus, Vibrio, D_4_Pasteurellaceae, Luteolibacter, Skermanella, Falsirhodobacter, Faecalibacterium, Agrococcus, Marinobacterium, D_2_OM190, D_4_Planococcaceae, Aggregatibacter, Aeromonas, Shinella, Chroococcidiopsis, Paraclostridium, Dolosigranulum, Bergeyella, D_1_Saccharibacteria, [Agitococcus] lubricus group, Paeniglutamicibacter, Aeromicrobium, Craurococcus, Nocardiopsis, Meiothermus, Hymenobacter, Exiguobacterium, Williamsia, Asticcacaulis, Serratia, Coprococcus 2, Marinomonas, Pectobacterium, Lautropia, [Eubacterium] yurii group, D_4_Xanthobacteraceae, Rickettsia, Schlesneria, Solobacterium, D_4_Clostridiales vadinBB60 group, Corynebacterium, Planktothrix, Kingella, Acidovorax, Gordonia, Parvimonas, D_4_Sandaracinaceae, Blautia, D_4_Myxococcaceae, D_2_Betaproteobacteria, Megamonas, Saccharopolyspora, Leucobacter, D_4_Oxalobacteraceae, Microvirga, D_4_Longimicrobiaceae, D_0_Bacteria;Ambiguous_taxa, Intestinibacter, D_4_Elev-16S-1332, Deefgea, Planomicrobium, Mogibacterium, Weissella, D_4_Cellulomonadaceae, Curtobacterium, Prevotella 9, D_4_Gemmatimonadaceae, Limnobacter, D_4_Ruminococcaceae, Chryseobacterium, Blastococcus, Haliangium, Enhydrobacter, Rubrivirga, Laceyella, Erwinia, Christensenellaceae R-7 group, Truepera, Quadrisphaera, Chthoniobacter, Oceanirhabdus, Zimmermannella, Flavisolibacter, D_4_YNPFFP1, Janibacter, Peptostreptococcus |
